# Ether lipid remodeling during neuronal differentiation prevents ferroptosis

**DOI:** 10.64898/2026.05.02.722203

**Authors:** Antonino Asaro, Gregor Pulikkotti Jose, Ilias Gkikas, Sarah Salame, Pauline Perné, Jean Andrea Maillat, Sylvia Ho, Océane Buvry, Lucile Fleuriot, Eva Bastida-Martínez, Jeremy Vicencio, Julián Cerón, Yuki Matsuzawa, Frédéric Brau, Julie Cazareth, Hiroshi Tsugawa, Delphine Debayle, Montserrat Elías-Arnanz, Howard Riezman, Giovanni D’Angelo, Takeshi Harayama

## Abstract

Ferroptosis is a form of cell death driven by iron-dependent lipid peroxidation, with specific lipid species playing key roles in modulating susceptibility. Among these, ether lipids have shown conflicting effects, being linked to both protection and sensitization. Here, we dissect the relationship between lipid structure and ferroptosis sensitivity and explain how ether lipids exert context-dependent effects. Ether lipids can promote ferroptosis through a metabolic bias towards the accumulation of polyunsaturated acyl chains and ethanolamine head groups, whereas this pro-ferroptotic tendency is counterbalanced by the anti-ferroptotic vinyl ether moiety introduced by plasmanylethanolamine desaturase 1. We show that this protective effect is critical for preventing ferroptosis in hiPSC-derived neurons, which accumulate otherwise pro-ferroptotic ether lipids during differentiation. This effect is not solely due to its antioxidant properties but also stems from the reprogramming of mitochondrial respiration. The lack of vinyl ether bonds leads to multiple mitochondrial defects, including increased mitochondrial reactive oxygen species (ROS), lower membrane potential, and abnormal cristae structures. These findings indicate that vinyl ether bonds in ether lipids offer dual ferroptosis resistance by scavenging ROS and minimizing its production at the mitochondrial level. The disruption of this system in *Caenorhabditis elegans* leads to iron-induced death and impaired motility. Thus, our study reveals ether lipid structural remodeling as a key regulator of ferroptosis sensitivity in neurons.

## INTRODUCTION

Ferroptosis is an iron-dependent form of cell death characterized by the accumulation of membrane phospholipid hydroperoxides, ultimately leading to membrane rupture and ionic dysregulation.^1^ Cellular susceptibility to ferroptosis is determined by the balance between peroxidizable lipid species, primarily polyunsaturated fatty acid (PUFA)-containing glycerophospholipids (GPLs), and antioxidant systems that suppress both the initiation and propagation of lipid peroxidation.^2^ Neurons are enriched in PUFA-containing GPLs,^3^ which makes it important to have intrinsic anti-ferroptotic mechanisms to maintain their survival. The breakdown of antioxidant systems in neural cells can trigger a ferroptotic cascade, which is a major driver of pathology and is often intertwined with neuroinflammation and iron dyshomeostasis.^4,5^ Indeed, dysregulated ferroptosis contributes to progressive neuronal loss in neurodegenerative diseases such as Alzheimer’s, Parkinson’s, amyotrophic lateral sclerosis, and multiple sclerosis.^4,5^

Ether GPLs are defined by a fatty alcohol linked at the *sn-*1 position via ether (for alkyl-GPLs) or vinyl ether linkages (for alkenyl-GPLs, also called plasmalogens) (Figure 1A). Ether GPLs have emerged as modulators of ferroptosis sensitivity.^6–8^ However, their role remains paradoxical. On one hand, several studies have identified that ether lipid biosynthesis promotes ferroptosis, likely because PUFA-rich ether GPLs are prone to peroxidation.^7,8^ On the other hand, a study showed that ether lipid deficiency sensitizes to ferroptosis,^6^ highlighting the need for more investigation on the relationship between ether lipids and ferroptosis. Indeed, plasmalogens, in which the vinyl ether bond is introduced by plasmanylethanolamine desaturase 1 (PEDS1),^9,10^ have long been associated with antioxidant functions and protection against oxidative stress, through their capacity to chemically scavenge oxidants.^11^

**Figure 1.**
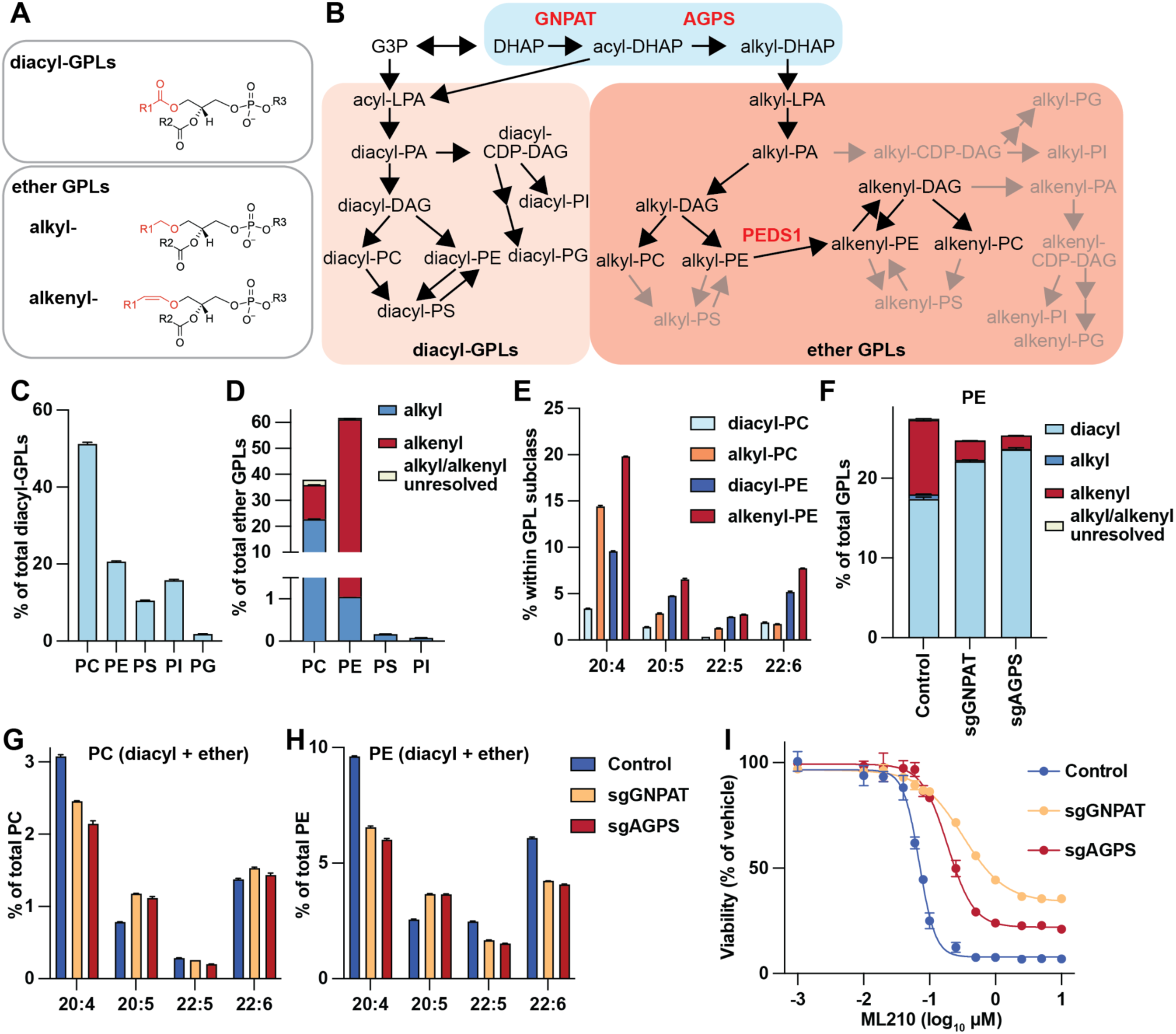
Ether lipids serve as a sink for PUFA and are pro-ferroptotic. (A) Structure of diacyl- and ether GPLs (see the region highlighted in red). (B) *De novo* synthesis of GPLs from glycerol 3-phosphate (G3P) or DHAP. PA: phosphatidic acid; DAG: diacylglycerol; PS: phosphatidylserine; PI: phosphatidylinositol; PG: phosphatidylglycerol. (C) Head group composition of diacyl-GPLs in HeLa cells. (D) Head group composition and linkage types of ether GPLs in HeLa cells. (E) Enrichment of various PUFAs in the indicated GPL subclasses. (F) Levels of PE subclasses (diacyl, alkyl, and alkenyl) in the indicated mutants. (G and H) Enrichment of major PUFAs in PC and PE, with all subclasses summed, across the indicated mutants. (I) Ferroptosis sensitivity of the indicated mutants. (C-I) Error bars are SEM of 4 biological replicates.

Ether GPL biosynthesis starts in peroxisomes from dihydroxyacetone phosphate (DHAP), which is converted into alkyl lysophosphatidic acid (alkyl-LPA) via multiple steps.^12^ Alkyl-LPA is metabolized into ether GPLs via reactions that are similar to those acting on its ester counterpart, acyl-LPA (Figure 1B).^12^ Nevertheless, ether GPLs have mainly choline or ethanolamine head groups and polyunsaturated *sn-*2 tails, which contrasts from diacyl-GPLs.^13^ This suggests that the ether linkage in alkyl-LPA dictates its metabolic fate differently from acyl-LPA, probably because some enzymes discriminate their substrates (a phenomenon we refer to as structure-guided metabolic bias).^14^

The contradictory findings related to ether lipid and ferroptosis raise key questions: Is the proposed pro-ferroptotic activity of ether lipids intrinsic to the ether (or vinyl ether) linkage itself, or does it stem from their metabolic bias by which they accumulate PUFA? To answer this question, it is critical to establish how ether lipid composition is metabolically regulated, and how ether lipid structure impacts ferroptosis. Another critical question related to ether lipids is what are the functions of the vinyl ether bond in plasmalogens beyond its proposed capacity to chemically scavenge oxidants.

In this study, we dissect the mechanisms of the metabolic bias of ether GPLs and study how their structure affects sensitivity to ferroptosis. We find that the accumulation of ethanolamine and PUFA tails in ether GPLs promote ferroptosis, which is counter-balanced by the introduction of a vinyl ether bond in plasmalogens. We show that this bias toward ethanolamine accumulation is enhanced during neuronal differentiation, making it critical for neurons to prevent ferroptosis by the introduction of the vinyl ether bond. We demonstrate that plasmalogens are not merely passive antioxidant reservoirs, but actively optimize mitochondrial respiration, reducing reactive oxygen species (ROS) production at its source. This mitochondrial adaptation is essential for suppressing lipid peroxidation and ferroptosis. Furthermore, we established the *in vivo* relevance of these findings in *C. elegans*, where knockout of the PEDS1 orthologue causes hypersensitivity to iron, leading to defects in survival and locomotion.

The results reveal multifaceted roles of plasmalogens in cells and *in vivo*. They clarify the role of plasmalogens in ferroptosis and uncover a previously unrecognized and evolutionary conserved metabolic safeguard that becomes hardwired during neuronal differentiation, offering potential therapeutic avenues to protect neurons from oxidative cell death.

## RESULTS

### The fatty acid composition of ether lipids determines ferroptosis sensitivity

We performed untargeted lipidomics of HeLa cells to obtain profiles of ether and diacyl-GPLs. Our approach discriminates alkyl- and alkenyl-GPLs from their retention time in LC-MS/MS, in addition to the annotations based on mass-to-charge ratio (m/z) and MS/MS fragmentations (similarly to a published strategy^15^). Ether GPLs have almost exclusively ethanolamine and choline head groups with the former being more abundant, in contrast to diacyl-GPLs that have choline, ethanolamine, inositol, serine, and glycerol (in order of abundance) as head groups (Figures 1C, 1D, and S1A). Ether phosphatidylcholine (PC) was more enriched in alkyl chains than alkenyl chains, while ether phosphatidylethanolamine (PE) had almost exclusively alkenyl chains (Figures 1D and S1A). Ether GPLs were richer in arachidonic acid (AA, 20:4) than their non-ether counterparts within the same lipid class, while PE was in general richer in PUFAs than PC (Figure 1E). Thus, despite sharing similar metabolic pathways with acyl-LPA (Figure 1B), the metabolism downstream of alkyl-LPA is highly biased towards ethanolamine and PUFA acquisition.

Glyceronephosphate *O-*acyltransferase (GNPAT) and alkylglycerone phosphate synthase (AGPS) are two peroxisomal enzymes involved in the initial steps of ether lipid biosynthesis (Figure 1B). To investigate the role of ether lipids in ferroptosis, we generated HeLa cell lines lacking GNPAT or AGPS (Figure S1B) and analyzed their lipidome and ferroptosis sensitivity. These mutant cells exhibited a strong reduction, but not a complete lack, in ether lipids (Figures 1F and S1C-S1E). Minor ether lipids with 20-carbon *sn-*1 fatty alcohols were unchanged in the mutants (Figure S1E). We found that the serum used for culture contains ether lysophosphatidylcholine (LPC) including 20-carbon ones, and that most of them are consumed in conditioned medium (Figure S1F). This suggests that ether LPC in serum is used by HeLa cells to make ether lipids, which compensates their loss in GNPAT and AGPS mutants.

Lipid tail analysis revealed that GNPAT and AGPS mutants have more PUFAs in diacyl-PE and diacyl-PC than in controls (Figures S1G and S1H), which partially compensated the decreases in GPL polyunsaturation due to the reductions in ether lipids. As a result, PC (including both diacyl and ether species) had similar PUFA levels in the mutants, while in PE the compensation was insufficient and PUFA levels were lower than in controls (Figures 1G and 1H). These results suggest that ether GPLs normally act as PUFA reservoirs, which increases the total mass of PUFA-containing GPLs in the form of PE.

To induce ferroptosis, we treated mutant HeLa cells with the glutathione peroxidase 4 (GPX4) inhibitor and ferroptosis inducer ML210.^16^ GPX4 inhibits ferroptosis by reducing lipid hydroperoxides into non-toxic lipid alcohols, using glutathione (GSH) as a cofactor.^17^ Both GNPAT and AGPS mutants were resistant to ferroptosis induced by ML210 (Figure 1I), confirming prior findings that ether lipid biosynthesis promotes ferroptotic sensitivity.^7,8^ A comparison between lipidomics data and ferroptosis sensitivity raised the possibility that the resistance of the mutants is due to decreased PUFA levels in PE (and not PC), rather than the changes in *sn-*1 linkages themselves (Figures 1G-1I).

Thus, we investigated how PUFA accumulation occurs in ether GPLs. A CoA-independent transacylase activity that transfers AA from diacyl-GPLs to ether lysophospholipids (ether lyso-GPLs) was first described in the 1980s^18^ (Figure 2A), yet the enzymes responsible remained unidentified at the onset of this study. Using the DepMap portal,^19^ which compiles genome-wide CRISPR-Cas9 screenings for essential genes of >1200 cell lines, we discovered that loss of TMEM164 leads to a selective growth disadvantage in cell lines that overlap with those that are sensitive to the loss of PEDS1 or lysophosphatidylcholine acyltransferase 3 (LPCAT3), an enzyme responsible for AA incorporation in GPLs (Figures 2A, 2B, and S1I). Being on the intersection of ether GPL metabolism and AA incorporation, we hypothesized that TMEM164 mediates the elusive transacylase activity.

**Figure 2.**
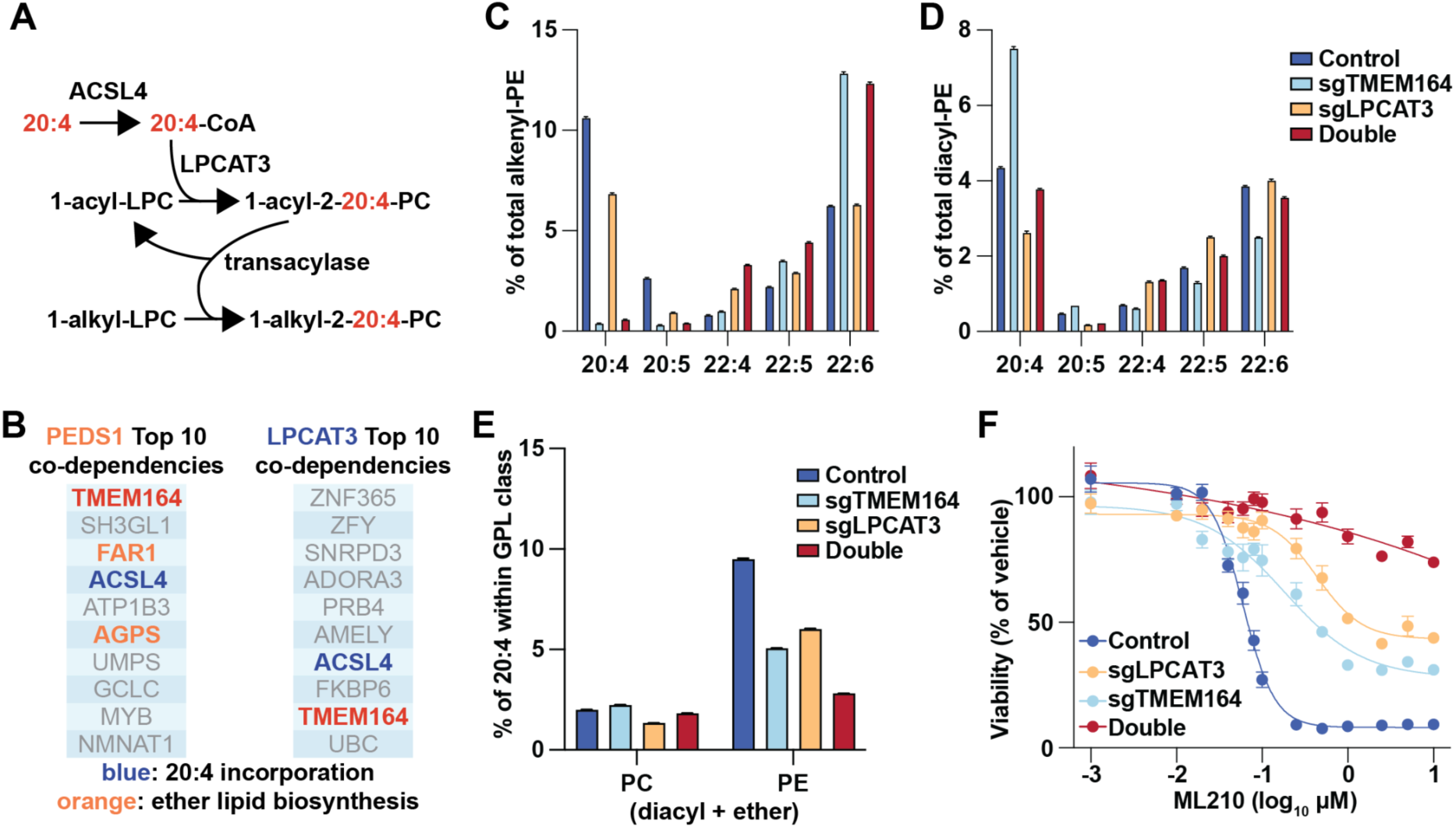
TMEM164 enriches AA in ether GPLs and renders them pro-ferroptotic. Genes that have essentiality profiles that correlate (positively or negatively) the most with PEDS1 and LPCAT3. Note the presence of TMEM164 in both. (C and D) Enrichment of abundant PUFAs in alkenyl-PE in diacyl-PE across the indicated mutants. (E) Enrichment of AA in PC and PE, with all subclasses summed, across the indicated mutants. (F) Ferroptosis sensitivity of the indicated mutants. (C-F) Error bars are SEM of 4 biological replicates.

To test this, we generated TMEM164 mutant HeLa cells and observed a marked reduction of AA in ether GPLs, accompanied by its accumulation in diacyl-GPLs, without large changes in the total levels of ether GPLs (Figures 2C, 2D, and S1J-S1M). The increases of AA in diacyl-PC compensated for the diminished levels in ether PC, while the compensation was incomplete for PE, leading to decreased AA levels in total PE (diacyl + ether PE) (Figure 2E). The results supported our hypothesis that TMEM164 is a transacylase that enriches AA in ether GPLs, leading to an increase in overall PUFA enrichment in PE. While this work was underway, a parallel study independently confirmed that TMEM164 functions as a CoA-independent transacylase, preferentially transferring AA from diacyl GPLs to ether lyso-GPLs.^20^ Immunofluorescence analysis localized TMEM164 to the endoplasmic reticulum, suggesting that AA-enrichment of ether GPLs occurs in this compartment (Figure S1N). Notably, TMEM164-deficient cells displayed resistance to ferroptosis, demonstrating that the accumulation of AA in ether GPLs is a key determinant of their pro-ferroptotic activity (Figure 2F). Interestingly, the enrichment of docosahexaenoic acid (DHA, 22:6) in alkenyl-PE was higher in TMEM164 mutants, which did not compensate for ferroptosis sensitivity (Figures 2C and 2F). This suggests that AA is more relevant than DHA for ferroptosis sensitivity, although the precise mechanisms for this tendency remain to be established.

**Figure S1.**
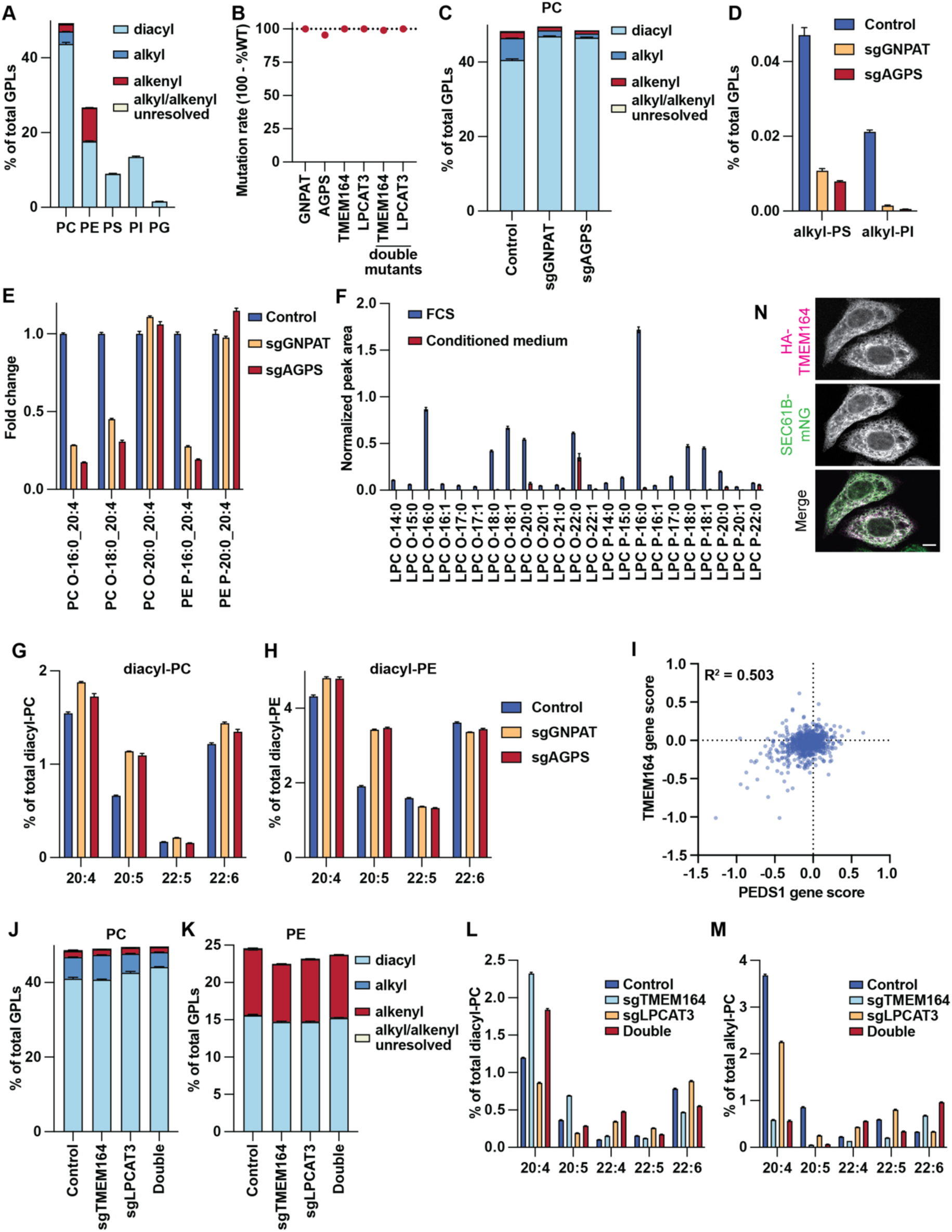
Lipidomic changes in mutant cells. (A) GPL profile of HeLa cells. (B) Mutation efficiencies in the indicated targets. (C) Levels of PC subclasses (diacyl, alkyl, and alkenyl) in the indicated mutants. (D) Changes in selected ether lipid species in the indicated mutants. O- and P-designate alkyl and alkenyl species, respectively. (F) Levels of various ether LPC species found in fetal calf serum (FCS) used to culture HeLa cells and in conditioned medium, in which FCS-derived lipids have been consumed. (G and H) Enrichment of major PUFAs in diacyl-PC and diacyl-PE across the indicated mutants. (I) Comparison of essentiality profiles between PEDS1 and TMEM164. Each dot represents a different cell line, with negative gene scores indicating that the depletion of the gene is lethal or gives a growth disadvantage. (J and K) Levels of subclasses of PC and PE in the indicated mutants. (L and M) Enrichment of abundant PUFAs in diacyl-PC and alkyl-PC across the indicated mutants. (N) Subcellular localization of TMEM164 identified by immunofluorescence. SEC61B-mNG is an endoplasmic reticulum marker fused to mNeonGreen. Scale bar = 5 µm. (A, C-H, and J-M) Error bars are SEM of 4 biological replicates.

We previously showed that LPCAT3 is responsible for AA enrichment in PC, PE, and phosphatidylserine (PS).^21^ We generated mutants of LPCAT3, either alone or in combination with TMEM164. LPCAT3 deficiency reduced AA levels in diacyl-PC, diacyl-PE, diacyl-PS, and ether GPLs (Figure 2C, 2D, S1B, S1L, and S1M). However, the decrease in AA-containing ether GPLs was less pronounced than in TMEM164 mutants. Thus, our results are in agreement with a model in which LPCAT3 enriches AA in some diacyl-GPLs, which are used as a substrate for TMEM164-dependent transacylation into ether GPLs. The dual deficiency of LPCAT3 and TMEM164 led to more pronounced decreases in AA levels than in single mutants, especially in total PE (Figure 2E). The deficiency of either enzyme was protective against ferroptosis, and the combinatory loss gave extremely strong protection (Figure 2F). By comparing lipidomics data and ferroptosis sensitivity in various mutants, we concluded that ferroptosis sensitivity scales the best with AA enrichment in PE, with the contribution not being restricted to ether PE. Thus, the sensitizing role of ether GPLs in ferroptosis sensitivity is likely a consequence of metabolic bias, which increases the overall polyunsaturation of PE.

### PEDS1 reduces the pro-ferroptotic potential of ether PE

To understand the mechanisms underlying the head group bias toward PE and PC in ether GPL synthesis, we examined how specific enzymatic preferences shape this distribution. In an unrelated project to study phosphatidylinositol (PI) metabolism, we generated HeLa cells ectopically expressing Saccharomyces cerevisiae CDP-diacylglycerol synthase 1 (ScCds1), and found that they accumulate alkyl-PI (Figure 3A). This finding indicates that ether PI can be synthesized if CDP-diacylglycerol synthase (CDS) activity is rewired. Of note, yeasts are devoid of ether lipids, thus no selective pressure would have led to an increased affinity of ScCds1 to ether substrates. Thus, we speculate that endogenous CDS enzymes in HeLa cells have evolved to exclude ether phosphatidic acid (PA) as a substrate (Figure 3B). This substrate rejection likely explains the general scarcity of ether PI in mammalian cells (Figure 1D).

**Figure 3.**
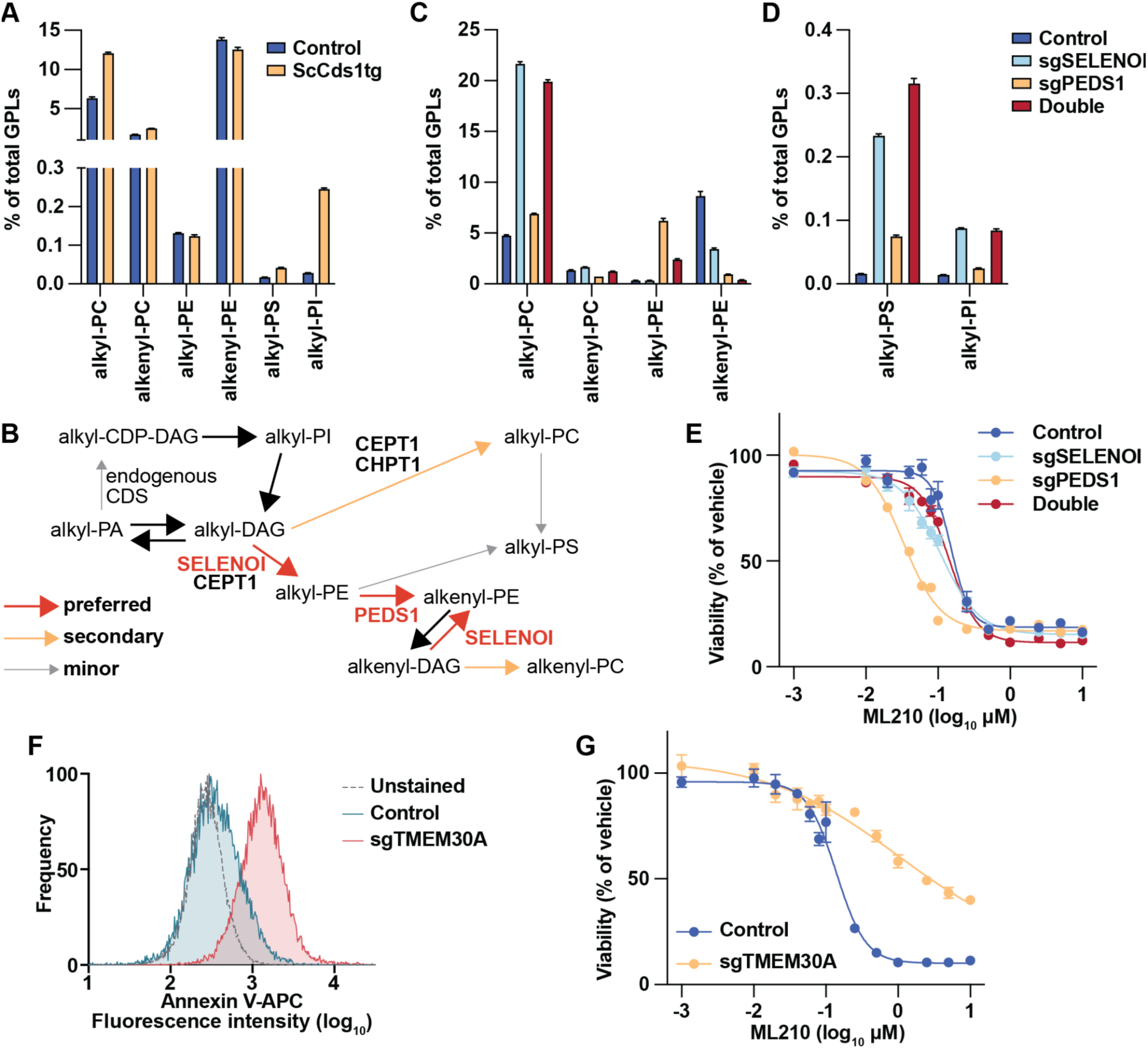
Head group, linkage, and asymmetry of GPLs affect ferroptosis. (A) Changes in ether lipid classes between control and ScCds1-overexpressing HeLa cells. (B) Metabolic pathways for ether lipids studied in this figure and Figure S2. The arrows are color coded depending on how biased the reactions are, with black arrows depicting reactions that were not directly addressed in this study. (C and D) Changes in ether lipid classes in the indicated mutants. (E) Ferroptosis sensitivity of the indicated mutants. (F) Analysis of cell surface PS exposure in control and TMEM30A mutants. (G) Ferroptosis sensitivity of control and TMEM30A mutants. (A, C-E, and G) Error bars are SEM of 4 biological replicates.

To further dissect head group specificity, we targeted enzymes that convert diacylglycerol (DAG) into PC or PE, namely CHPT1, CEPT1, and SELENOI (selenoprotein I) (Figures S2A and S2B). CHPT1 catalyzes PC synthesis, while CEPT1 produces both PC and PE (Figure S2A). SELENOI is also called ethanolamine phosphotransferase 1 (EPT1) and synthesizes PE with preference for ether species (Figures 3B and S2A).^22^ CHPT1 and CEPT1 mutant cells did not have drastic changes in ether GPL levels, with only statistically non-significant decreases in alkyl-PC observed (Figure S2C). CEPT1 mutants had decreases in PS and increases in PI (Figure S2C). Thus, CHPT1 and CEPT1 do not have major contributions on the metabolic biases of ether lipids. In contrast, SELENOI mutants showed a marked reduction in alkenyl-PE and a striking compensatory increase in alkyl-PC (Figure 3C), which agrees with a previous study.^22^ Notably, alkyl-PS and alkyl-PI levels were also elevated (Figure 3D), showing that SELENOI has a strong ability to sequester alkyl substrates and prevent their use for the synthesis of other ether GPLs.

In parallel, we disrupted PEDS1, the desaturase that converts alkyl-PE into alkenyl-PE (plasmalogen PE) (Figures 3B and S2B).^9,10^ The mutants accumulated alkyl-PE with only minimal shifts in alkyl-PC (Figure 3C). Intriguingly, alkyl-PS was also increased, suggesting that phosphatidylserine synthase 2 (PSS2), which converts PE to PS, weakly accepts alkyl-PE but not alkenyl-PE as a substrate (Figure 3B).

**Figure S2.**
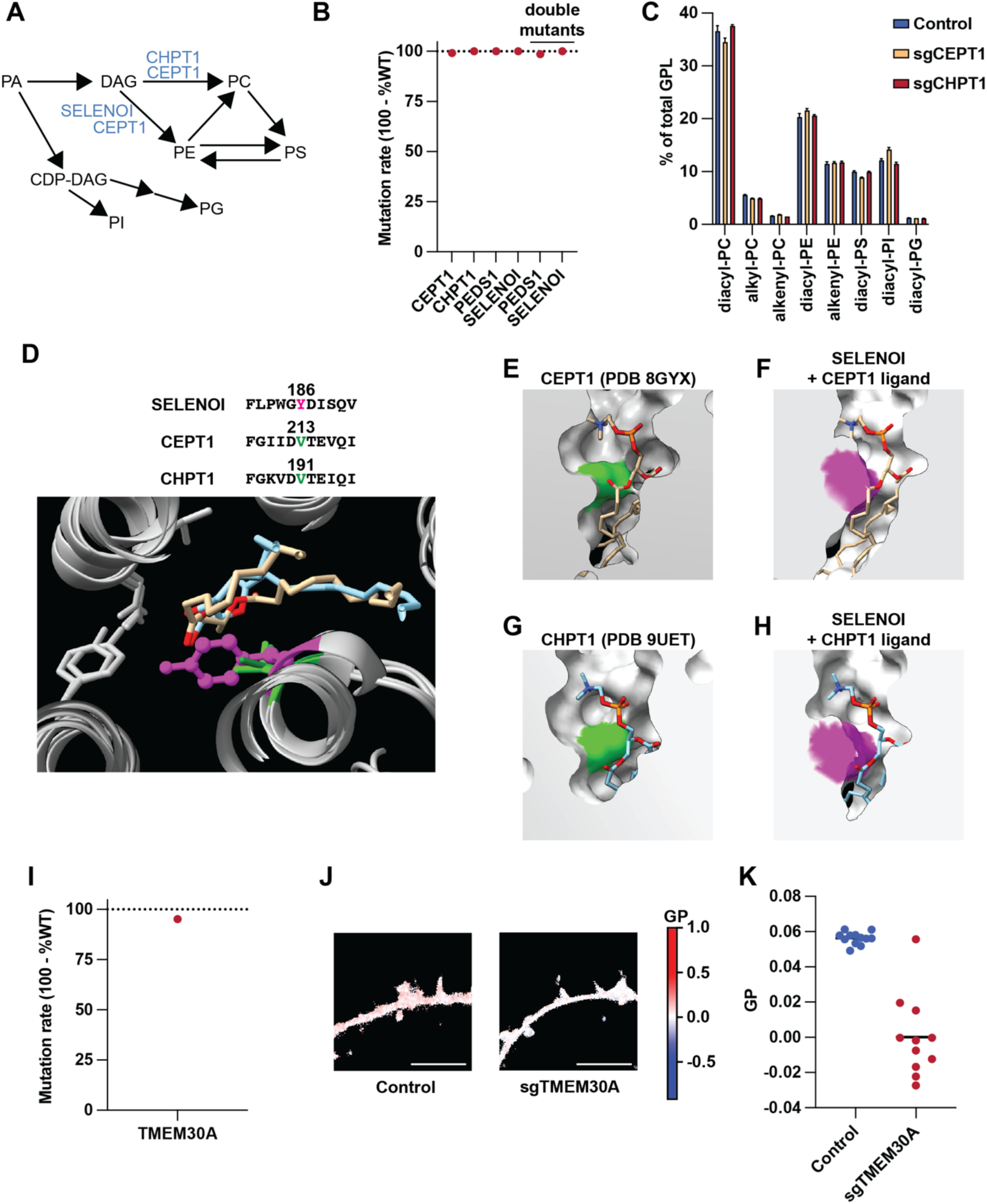
Characterization of PEDS1, SELENOI, and TMEM30A mutants. (A) Enzymatic steps catalyzed by CEPT1, CHPT1, and SELENOI. (B) Mutation rates in the target genes of the indicated mutants. (C) Changes in GPL subclasses in the indicated mutants. Error bars are SEM of 4 biological replicates. (D) Comparison of structures between SELENOI (prediction), CEPT1, and CHPT1, at regions close to the *sn-*1 linkages of substrates. (E-H) Comparison of substrate binding pocket shapes, with the substrates experimentally determined structures positioned in the equivalent position of predicted SELENOI structure. (I) Mutation efficiency of TMEM30A. (J and K) Analysis of lipid packing as generalized polarization values of Nile Red 12S in control and TMEM30A mutants. Scale bar = 5 µm.

Together, these results reveal that the head group bias in ether GPLs arises from three key enzymatic features: (1) SELENOI’s high specificity for ether-linked intermediates; (2) exclusion of ether PA by CDP-diacylglycerol synthase, which prevents ether PI formation; (3) limited ability of PS synthases to use ether substrates. This coordinated enzymatic architecture ensures the channeling of ether lipids predominantly toward PE and to a lesser extent PC (Figure 3B).

To gain structural hints for the mechanisms of ether lipid metabolic bias, we compared the predicted structure of SELENOI to experimental ones of CEPT1 and CHPT1, both having PC in their substrate binding pockets.^23,24^ SELENOI structure was not available in AlphaFold Protein Structure Database in full length, thus we used AlphaFold 3^25^ to generate a predicted structure of SELENOI with selenocysteine 387 replaced with a cysteine. The ester linkage at the *sn-*1 position of PC is proximal to valine 213 of CEPT1 or valine 191 of CHPT1 (Figure S2D). The analogous position in SELENOI is occupied by a bulkier tyrosine 186 (Figure S2D). This makes the substrate-binding pocket in SELENOI narrower at the region close to the *sn-*1 linkage, which would lead to steric clashes with the carbonyl oxygen of the ester linkage, at least if the substrate adopts the same configuration as in CEPT1 or CHPT1 (Figures S2E-S2H). Our results predict that the ether substrate preference of SELENOI is related to the shape of its substrate-binding pocket, which accommodates better *sn-*1 ether linkages that lack the carbonyl oxygen.

We next assessed the sensitivity of SELENOI and PEDS1 mutant HeLa cells to ferroptosis. Crucially, PEDS1 mutants showed increased susceptibility to ferroptosis, while SELENOI mutants displayed no significant change (Figure 3E). This suggests that an accumulation of alkyl-PE, but not alkyl-PC, enhances ferroptotic sensitivity (compare Figures 3C and 3E). To test this, we generated SELENOI/PEDS1 double mutant cells (Figure S2B) and compared them to PEDS1 single mutants. Both mutants have reductions in alkenyl-PE, with the increase in alkyl-PE being less pronounced in the double mutants (Figure 3C). Remarkably, the double mutants were more resistant to ferroptosis than PEDS1 single mutants, confirming that the ethanolamine head group in alkyl-GPLs, rather than the ether linkage itself, confers ferroptotic sensitivity (Figure 3E). Our results also show that PEDS1 is anti-ferroptotic only when SELENOI is present, as SELENOI single mutants and SELENOI/PEDS1 double mutants have similar ferroptotic sensitivity (Figure 3E).

We hypothesized that the pro-ferroptotic effect of ethanolamine head groups may arise from their preferential enrichment in the plasma membrane cytosolic leaflet, mediated by flippases.^26^ This positioning could make them more accessible to lipid peroxidation propagating from intracellular sources. To test this, we disrupted lipid asymmetry by mutating TMEM30A (Figure S2I), which encodes CDC50A, the essential subunit of P4-type ATPase flippases.^26^ As expected, TMEM30A mutant cells showed elevated PS levels on the outer plasma membrane (PM) leaflet, indicating impaired lipid asymmetry (Figure 3F).

To test if impaired lipid asymmetry is accompanied with altered PM physical properties, we measured the packing of the outer leaflet using the polarity-sensitive dye Nile Red 12S.^27^ This revealed that the PM outer leaflet is less ordered in TMEM30A mutants, consistent with the exposure of polyunsaturated GPLs that are normally present in the inner leaflet (Figures S2J and S2K).^28^ Thus, TMEM30A mutants have impaired asymmetry in GPL composition and unsaturation. These mutants were also more resistant to ferroptosis, supporting our hypothesis that polyunsaturated lipids in the inner leaflet sensitize cells to ferroptosis (Figure 3G).

Collectively, our results demonstrate that the apparent pro-ferroptotic potential of ether GPLs stems from two converging metabolic biases: (1) their enrichment in AA mediated by TMEM164, and (2) their preferential acquisition of ethanolamine head groups by SELENOI. PEDS1 counterbalances excessive ferroptosis when this pro-ferroptotic combination between ethanolamine head groups and polyunsaturated tails accumulates, by introducing vinyl ether linkages that have important anti-ferroptotic functions.

### Ether GPL remodeling during neural differentiation protects against ferroptosis

Next, we investigated the physiological relevance of PEDS1 in counterbalancing the ferroptosis sensitization driven by SELENOI-dependent ether PE synthesis. We focused our study on human neurons as they are highly vulnerable to oxidative damage^29^, while mutations in SELENOI and PEDS1 cause neurodevelopmental diseases, showing the importance of plasmalogen synthesis in neurons.^22,30,31^ Therefore, we tested whether human neurons exhibit ether PE and PUFA-rich lipid profiles and whether PEDS1 protects them against ferroptosis.

We employed neurogenin 2 (NGN2)-mediated conversion of human induced pluripotent stem cells (iPSCs) into neurons (iN). Lipids were extracted from iPSCs, early neuronal precursors (day-5, d5), and mature neurons (day-14, d14), followed by LC-MS-based lipidomics (Figure 4A). We observed distinct lipidomic profiles associated with different stages of neuronal differentiation (Figures 4B-4F and S3). We found a systematic decrease in PS, phosphatidylglycerol, and sphingolipids, along with an increase in PE across neuronal differentiation (Figures 4C, 4D, 4F, and S3).

**Figure 4.**
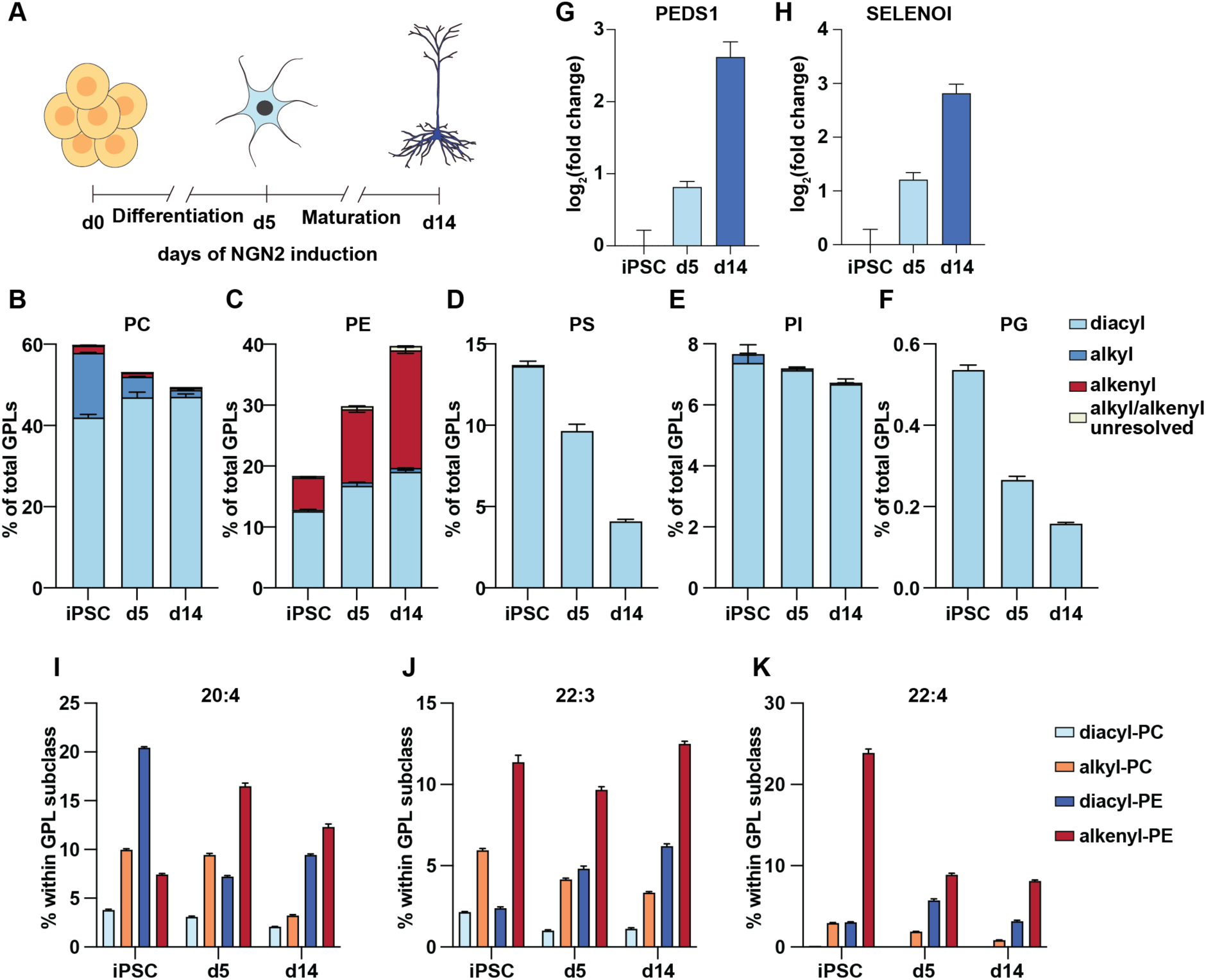
Ether GPLs are remodeled during neuronal differentiation. (A) Schematic overview of the NGN2-induced neuronal differentiation from iPSCs to day-5 precursors (d5) and day-14 neurons (d14). (B-F) Head group composition and *sn-*1 linkages of GPLs in iPSCs, d5, and d14 neurons. (G and H) qPCR analysis of PEDS1 and SELENOI expression during neuronal differentiation. (I-K) Enrichment of the indicated PUFAs within GPL subclasses during neuronal differentiation. (B-K) Error bars are SEM of 6 biological replicates.

Notably, the increase in PE was mainly a consequence of drastic remodeling that occurs in ether GPLs during differentiation. Alkyl-PC is enriched in iPSCs, while alkenyl-PE becomes predominant in d14 mature neurons (Figures 4B and 4C). Consistent with this shift, PEDS1and SELENOI mRNA levels increased in mature neurons compared to early-stage neurons (Figures 4G and 4H). The analysis of acyl chain enrichment in GPL subclasses revealed that alkenyl-PE becomes the major pool for PUFA esterification after neuronal differentiation (Figures 4I-4K), which is a situation where anti-ferroptotic vinyl ether bonds are important.

When we mutated PEDS1 in iPSCs and converted them into iN (Figure S4A), the mutant lacked alkenyl-GPLs with strong increases in alkyl-PE (Figures S4B and S4C). Despite this major shift in ether lipid composition, PEDS1-deficient cells differentiated normally into neurons, as reflected by both transcriptomic profiles (Figure S4D) and stage-specific marker expression across iPSCs, d5 precursors, and d14 neurons (Figure S4E).

**Figure S3.**
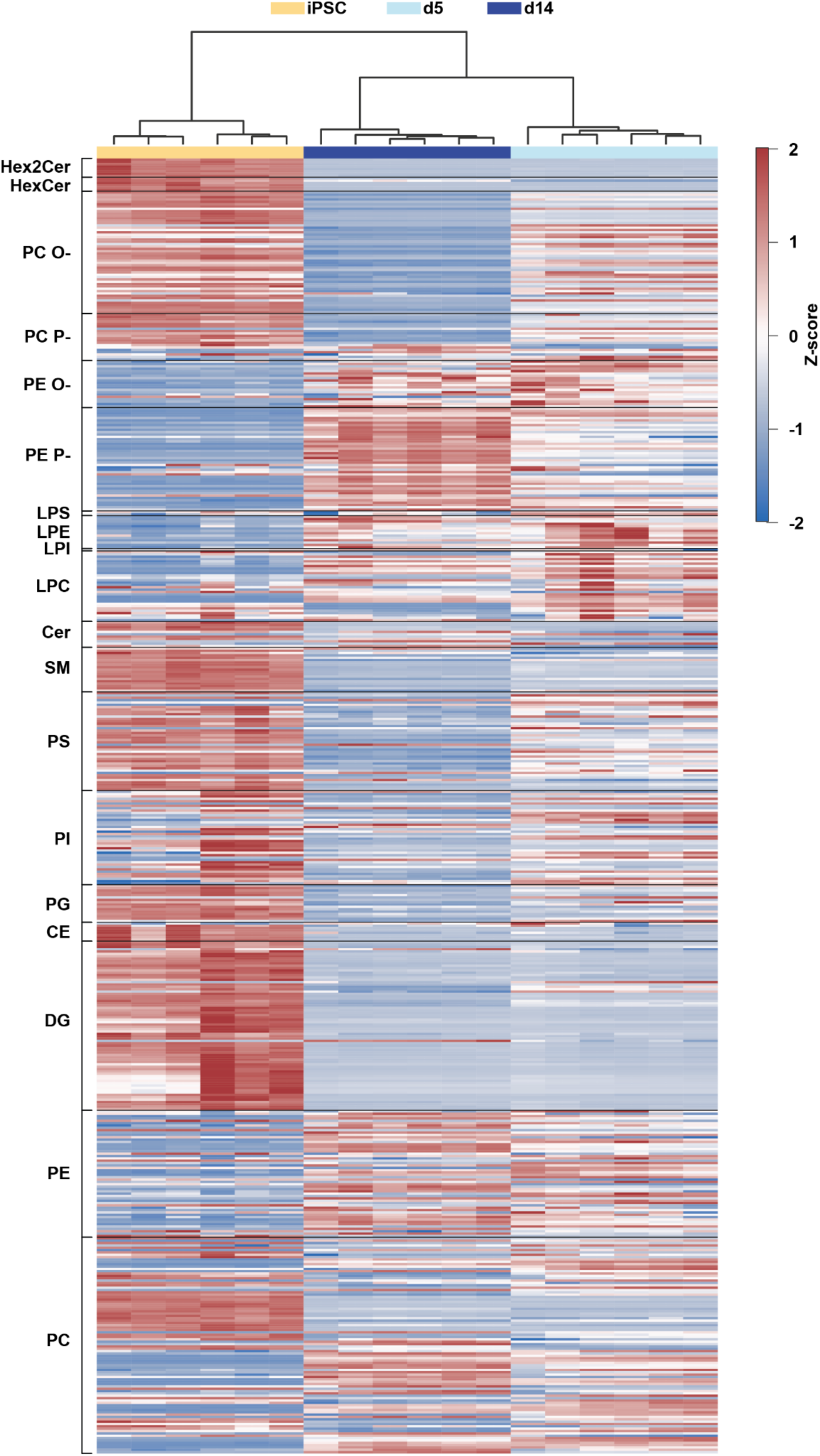
Lipid remodeling during neuronal differentiation. Heatmap of lipid species labelled by lipid class/subclass and clustered by condition across neuronal differentiation.

Standard media used for differentiating iPSCs into neurons contain substantial amounts of antioxidant additives. To mimic mild oxidative conditions, we transferred iPSC-derived neurons to a medium lacking these supplements.^32^ While neurons derived from control iPSCs survived in antioxidant-free medium for several days, PEDS1 mutant neurons succumbed rapidly (Figures 5A and 5B). This effect was not rescued by the pan-caspase inhibitor Z-VAD-FMK^33^, indicating that apoptosis was not the primary cause (Figures 5C and 5D). In contrast, viability was completely restored by the ferroptosis inhibitor ferrostatin-1^1^ (Figures 5E and 5F). Consistent with these findings, Liperfluo staining, an indicator of lipid peroxidation^34^, revealed increased lipid peroxidation in PEDS1 mutant neurons (Figure 5G). We then assessed ROS levels. As expected, antioxidant withdrawal increased endogenous ROS in all conditions, but neurons lacking plasmalogens exhibited significantly higher levels than controls (Figure 5H). Thus, PEDS1 has antioxidant and anti-ferroptotic roles in neurons, demonstrating the importance of ether lipid remodeling during differentiation.

**Figure 5.**
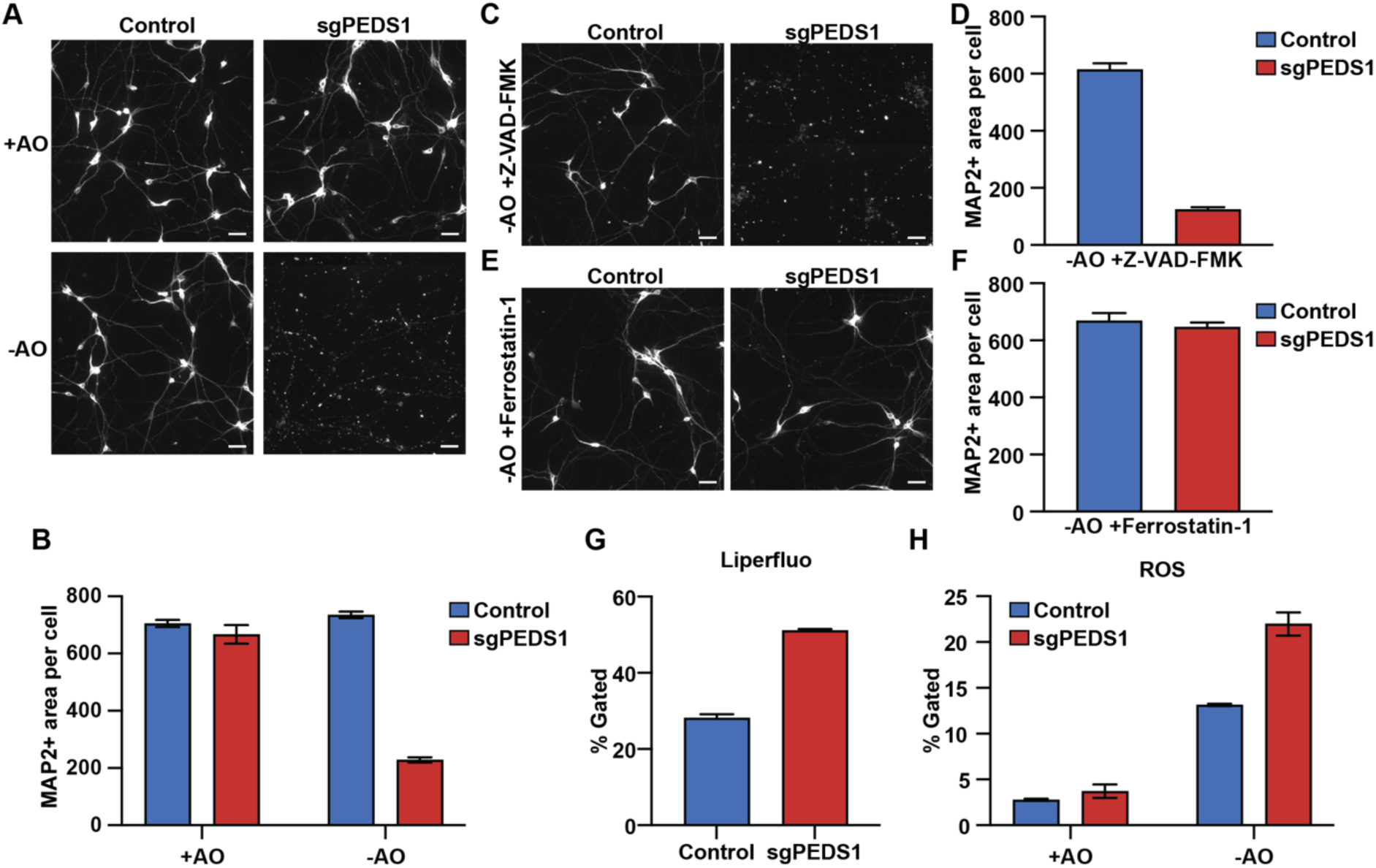
Absence of plasmalogen lipids sensitizes neurons to ferroptosis. (A and B) Immunostaining and quantification of MAP2-positive dendritic area in control and PEDS1 mutant neurons grown for 3 days in antioxidant-containing medium (+AO) or in medium lacking antioxidants (−AO). (C-F) Rescue experiments showing MAP2-positive dendritic area and quantification following treatment with the apoptosis inhibitor Z-VAD-FMK or the ferroptosis inhibitor Ferrostatin-1 in neurons cultured under −AO conditions. (G) Flow cytometry-based quantification of ferroptosis using the lipid peroxidation probe Liperfluo in control and PEDS1 mutant neurons cultured under −AO conditions. (H) Flow cytometry-based quantification of intracellular reactive oxygen species (ROS) in control and PEDS1 KO neurons in +AO or −AO medium. Scale bar = 50 µm. (B, D, and F-G) Error bars are SEM of 3 replicates.

**Figure S4.**
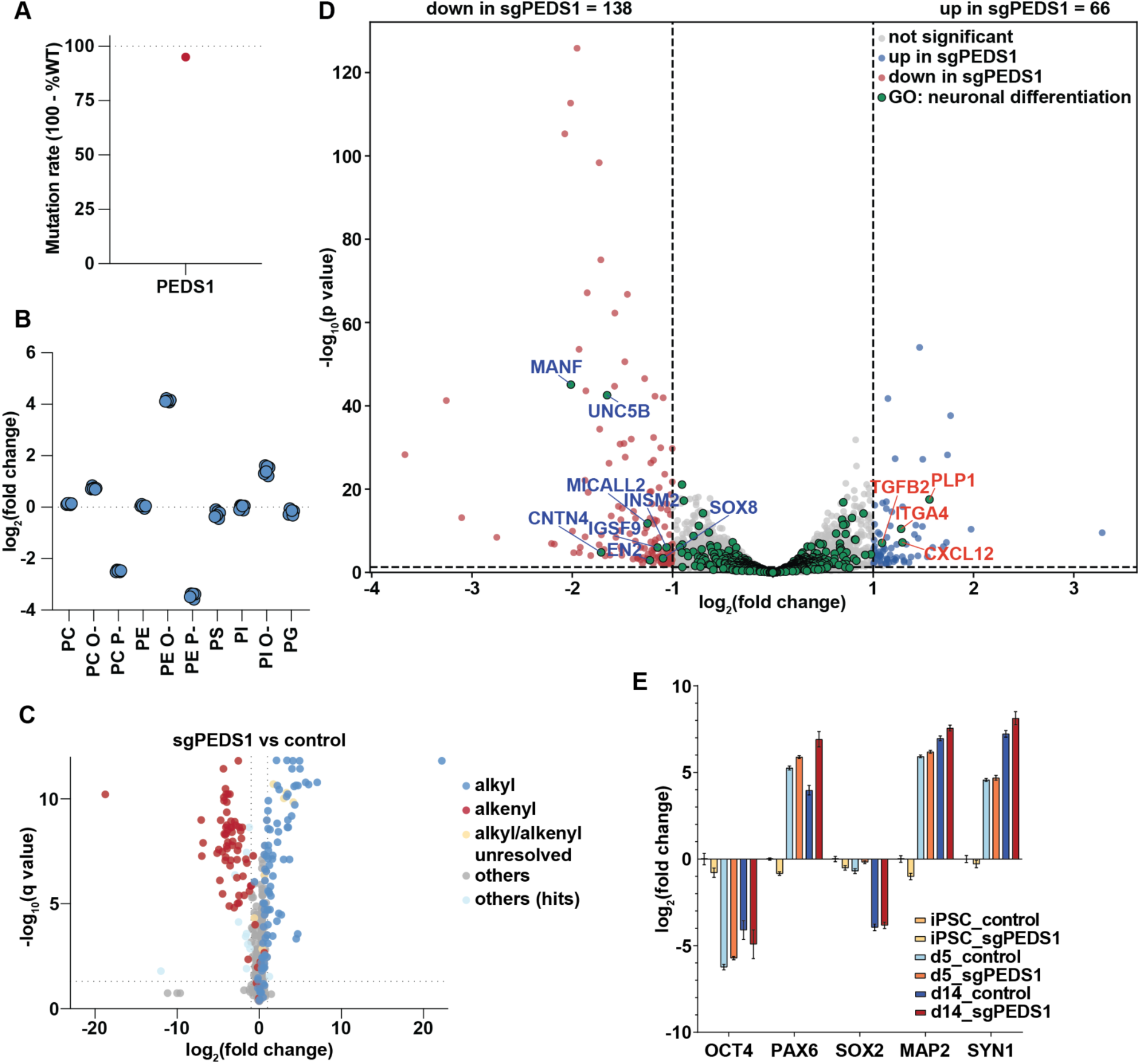
Characterization of PEDS1 KO neuronal cells. (A) TIDE analysis (Tracking of Indels by DEcomposition) validating CRISPR/Cas9-based disruption of PEDS1 in iPSC cells. (B) Fold-change analysis of lipid subclass composition from lipidomic profiling of PEDS1 mutant and control neurons. (C) Volcano plot of lipidomic analysis comparing PEDS1 mutant and control neurons, highlighting ether lipid subclasses in different colors. (D) Volcano plot of RNA-seq data from mature neurons comparing control and PEDS1 mutants, highlighting genes associated with neuronal differentiation in green. (E) qPCR analysis of pluripotency and neuronal differentiation markers in iPSCs, early neuronal precursors (day-5, d5), and mature neurons (day-14, d14) with or without PEDS1. Error bars are SEM of 6 biological replicates.

### Plasmalogens regulate mitochondrial function and membrane architecture

Since lipid peroxidation propagates from internal organelles to the plasma membrane during ferroptosis,^38^ we examined the localization of signals from the lipid peroxidation sensor C11-BODIPY. Shortly after antioxidant withdrawal, oxidized C11-BODIPY^39^ signals often colocalized with mitochondria (Figure S5A). Furthermore, PEDS1 mutant neurons exhibited significantly elevated mitochondrial superoxide levels (Figures 6A and 6B). This mitochondrial dysfunction appears causal to cell death, as pretreatment with the mitochondria-targeted antioxidant MitoQ^40^ fully restored viability under deprivation of antioxidants (Figures 6C and 6D).

**Figure 6.**
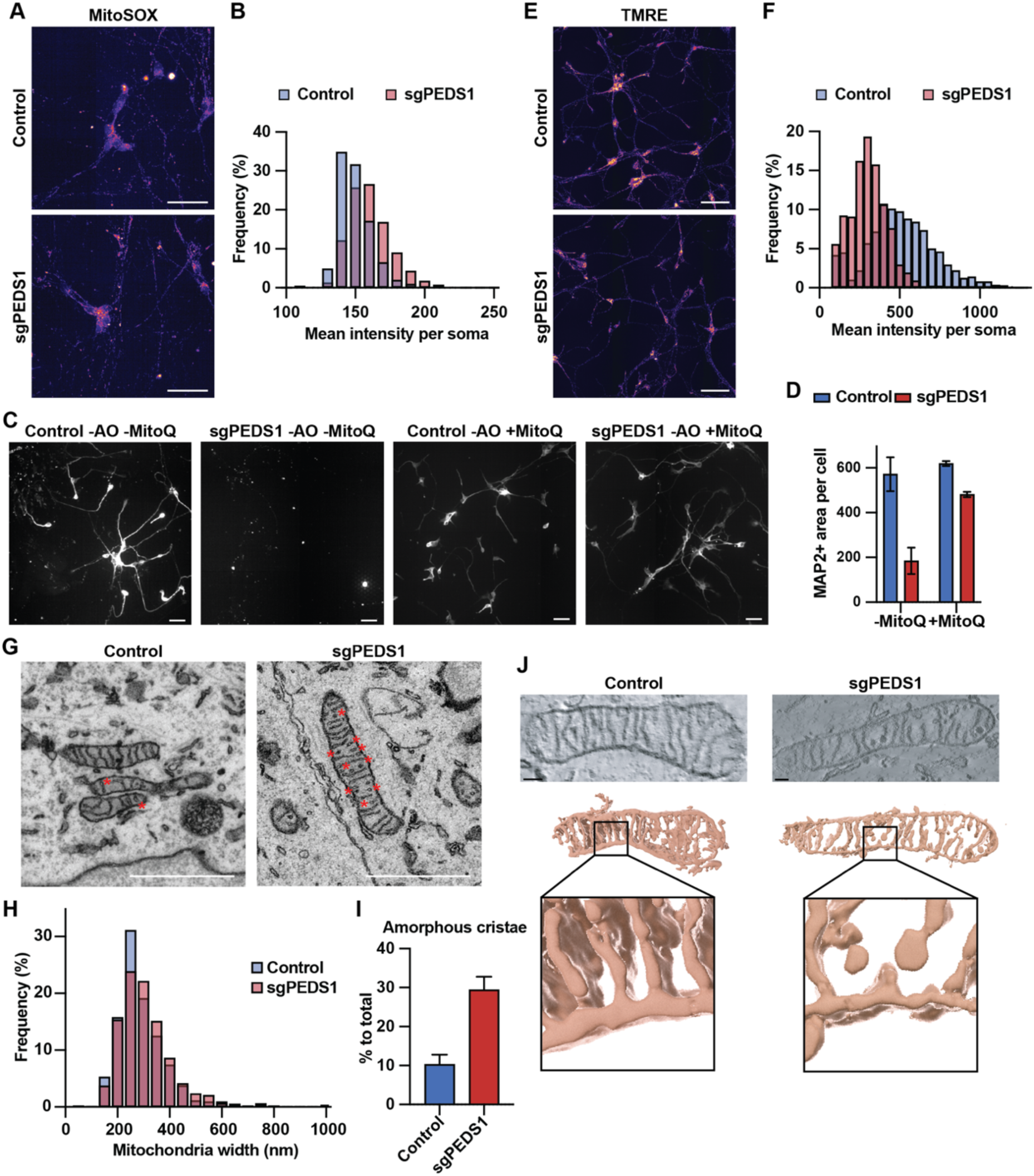
Plasmalogen loss impairs mitochondrial functions and morphology. (A) Staining of mitochondrial superoxide with MitoSOX and (B) frequency distributions of MitoSox mean fluorescence intensity of day-14 control and PEDS1 mutant neurons. (C and D) Map2 staining and quantification of MAP2-positive area of Control and PEDS1 mutant neurons cultured for 3 days in medium without antioxidants (−AO) with or without pretreatment with MitoQ. Error bars are SEM of 3 replicates. (E) Staining of mitochondrial membrane potential using TMRE dye and (F) frequency distributions of TMRE mean fluorescence intensity of day-14 control and PEDS1 mutant neurons. (G) Representative electron microscopy images of control and PEDS1 mutant neurons. (H) Frequency of individual mitochondrial width found in electron micrographs. (I) Percentage of mitochondria exhibiting “amorphous” cristae architecture. Error bars are SEM of 47 control and 43 sgPEDS1 replicates. (J) Representative tomography electron microscopy images of control and PEDS1 mutant neurons with 3D reconstruction, highlighting bulbous protrusion of cristae in PEDS1 mutants. Scale bars = 50 µm for fluorescence microscopy, 1 µm for electron microscopy, and 0.1 µm for cryo-electron tomography.

Neurons depend on mitochondrial respiration to meet their high energy needs, yet mitochondria generate ROS during oxidative phosphorylation.^41^ We thus explored whether plasmalogen loss impacts mitochondrial functions and bioenergetics. We first stained mitochondrial proteins to investigate their abundance, and found no differences between control and PEDS1 mutant neurons (Figure S5B). We then profiled the proteome of crude mitochondrial fractions from PEDS1 mutant neurons and observed no dysregulation of mitochondrial proteins (Figure S5C). Thus, numbers and protein contents of mitochondria seemed unaffected by PEDS1 loss. On the other hand, oxygen Consumption Rate (OCR) was slightly elevated in PEDS1 mutant neurons (Figures S5D and S5E). This metabolic signature was conserved in PEDS1 mutant HeLa cells (Figure S5F), showing that PEDS1 reduces OCR irrespectively of cell type. Notably, these metabolic assessments (OCR and mitochondrial ROS) were performed under standard culture conditions containing antioxidants, indicating an intrinsic metabolic reprogramming independent of oxidative stress.

Given the increased oxygen consumption and ROS production in PEDS1 mutant neurons, we investigated whether structural defects in the electron transport chain were responsible. Although Complexes I and III are primary sites of ROS generation, native gel electrophoresis revealed no changes in these complexes (Figures S5G-S5I). Also, we found that higher respiration did not result in increased ATP production (Figure S5G), suggesting mitochondrial uncoupling. We confirmed this hypothesis using TMRE staining,^42^ which demonstrated a significant reduction in mitochondrial membrane potential in PEDS1 mutant neurons compared to controls (Figures 6E and 6F). Thus, the absence of plasmalogens leads to a futile cycle of high respiration with associated ROS production.

**Figure S5.**
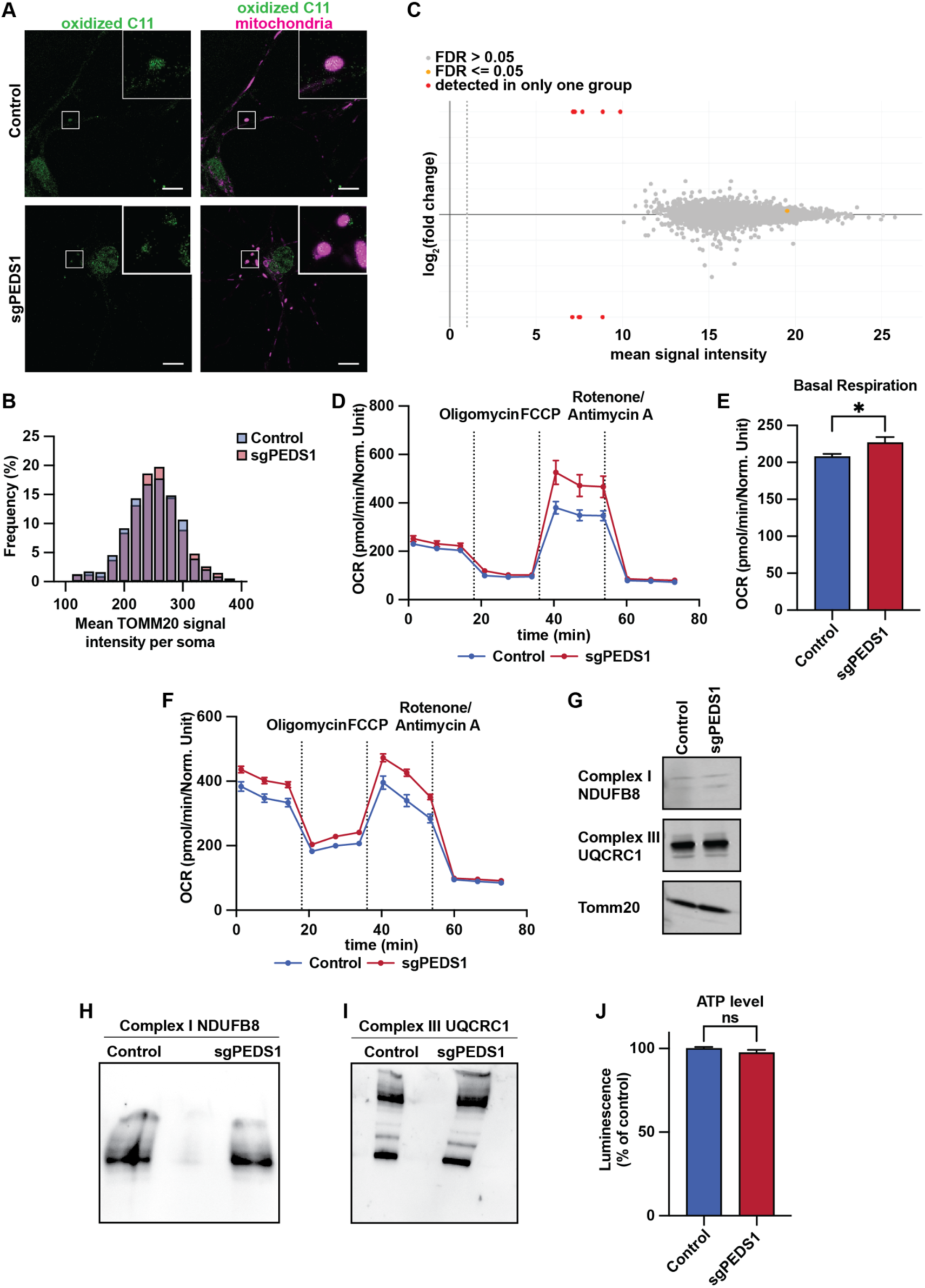
Plasmalogen deficiency affects mitochondrial functions. (A) Confocal imaging of oxidized C11-BODIPY (green) and MitoTracker (magenta) in control and PEDS1 mutant neurons after 8 h in antioxidant-free medium. Colocalization indicates that early lipid peroxidation occurs at mitochondria. Scale bar = 10 µm. (B) Frequency distributions of the mean fluorescence intensity of the mitochondrial marker TOMM20 in day-14 control and PEDS1 mutant neurons. (C) Proteomic profiling of crude mitochondrial fractions from control and PEDS1 mutant neurons. MA plot shows no consistently dysregulated mitochondrial pathways following plasmalogen loss. (D-F) Oxygen consumption rate (OCR) in (D and E) day-14 neurons (day-14) and (F) HeLa cells measured using the Seahorse XF Mito Stress Test. The indicated drugs were sequentially injected to assess OCR in various mitochondrial respiratory states. (G-I) Western blot of respiratory chain complex (Complexes I and III) proteins in control and PEDS1 mutant neurons, separated in denaturing gels (G) or by Blue Native PAGE (H and I). (J) Cellular ATP levels measured by CellTiter-Glo. (D-F and J) Error bars are SEM of (D) 10, (E) 20, (F) 20, and (J) 10 replicates.

To understand the structural basis of this mitochondrial uncoupling, we examined cristae architecture. Cristae are highly curved inner membrane folds supported by specialized lipids;^43^ their geometry is critical for maintaining respiratory efficiency.^44^ Electron microscopy revealed that while overall mitochondrial size and gross morphology were comparable between genotypes, cristae architecture was altered in PEDS1 mutant neurons (Figures 6G and 6H). Control mitochondria exhibited well-defined lamellar cristae, whereas PEDS1 mutant mitochondria displayed frequently an irregular organization that appeared ring-like in sections, which we tentatively named “amorphous cristae” (Figures 6G and 6I). Because such discontinuities can arise from different underlying crista morphologies, we used electron tomography to further resolve crista organization. Electron tomography supported altered crista organization in PEDS1 mutant neurons, revealing crista membranes with irregular geometry and bulbous protrusions rather than continuous lamellar folds (Figure 6J).

Collectively, these data indicate that plasmalogens contribute to mitochondrial membrane structure. Their loss disturbs mitochondrial function and enhances ROS production even under standard antioxidant-containing culture conditions. This underscores an intrinsic requirement for plasmalogens in maintaining mitochondrial efficiency and redox balance.

### Plasmalogen Deficiency Exacerbates Ferroptosis-Driven Iron Toxicity in C. elegans

To examine the role of plasmalogens *in vivo*, we employed ferric ammonium citrate (FAC) to overload *Caenorhabditis elegans* with iron as a pro-ferroptotic insult.^45^ Wild-type (wt) worms displayed a concentration-dependent decline in survival upon FAC exposure (Figure 7A). To determine whether plasmalogens play any role in this process, we used CRISPR-Cas genome-editing to generate a mutant of the worm orthologue of PEDS1. As a result, we produced a *C. elegans* strain harboring the deletion allele *tmem-189(cer184)*, hereafter referred to as *tmem-189(ko)* (see Materials and Methods). The *tmem-189(ko)* strain was devoid of plasmalogens and overproduced alkyl-GPLs, as expected (Figures S6A and S6B). *tmem-189(ko)* worms were significantly more vulnerable to FAC treatment, exhibiting mortality at FAC concentrations that were only mildly toxic to wt worms (Figures 7B and 7C).

**Figure 7.**
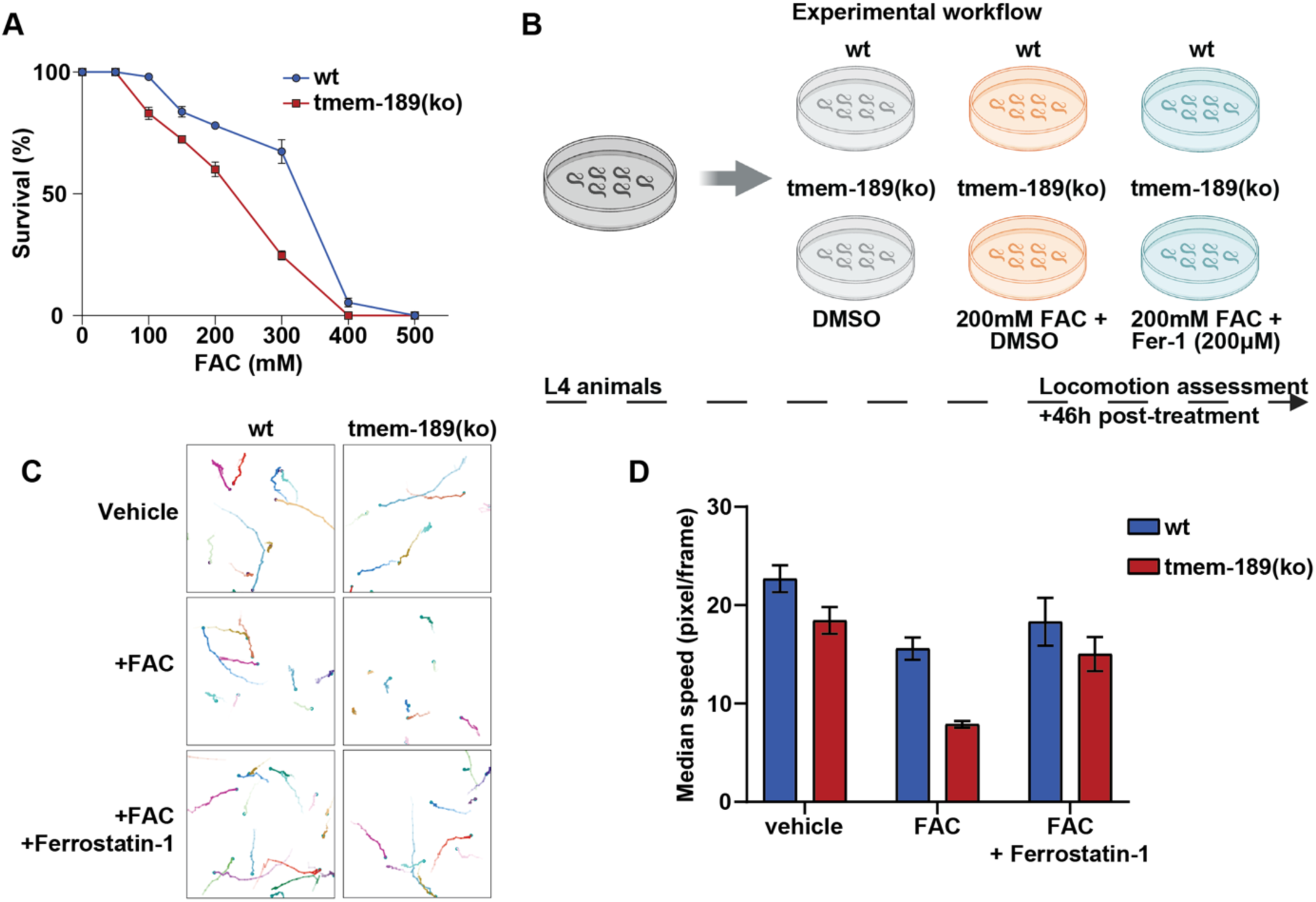
Plasmalogen deficiency increases sensitivity to iron-induced toxicity in *C. elegans*. (A) Survival analysis of wild-type (*wt*) and *tmem-189* mutant (ko) *C. elegans* exposed to increasing concentrati ns of ferric ammonium citrate (FAC). (B) Schematic experimental workflow of animal treatment for locomotion analysis. (C) Locomotor performance and (D) quantification of wild-type and *tmem-189* mutant animals exposed to sublethal FAC concentrations and rescue with ferrostatin-1 (Fer-1). Error bars are SEM of 18-39 replicates.

**Figure S6.**
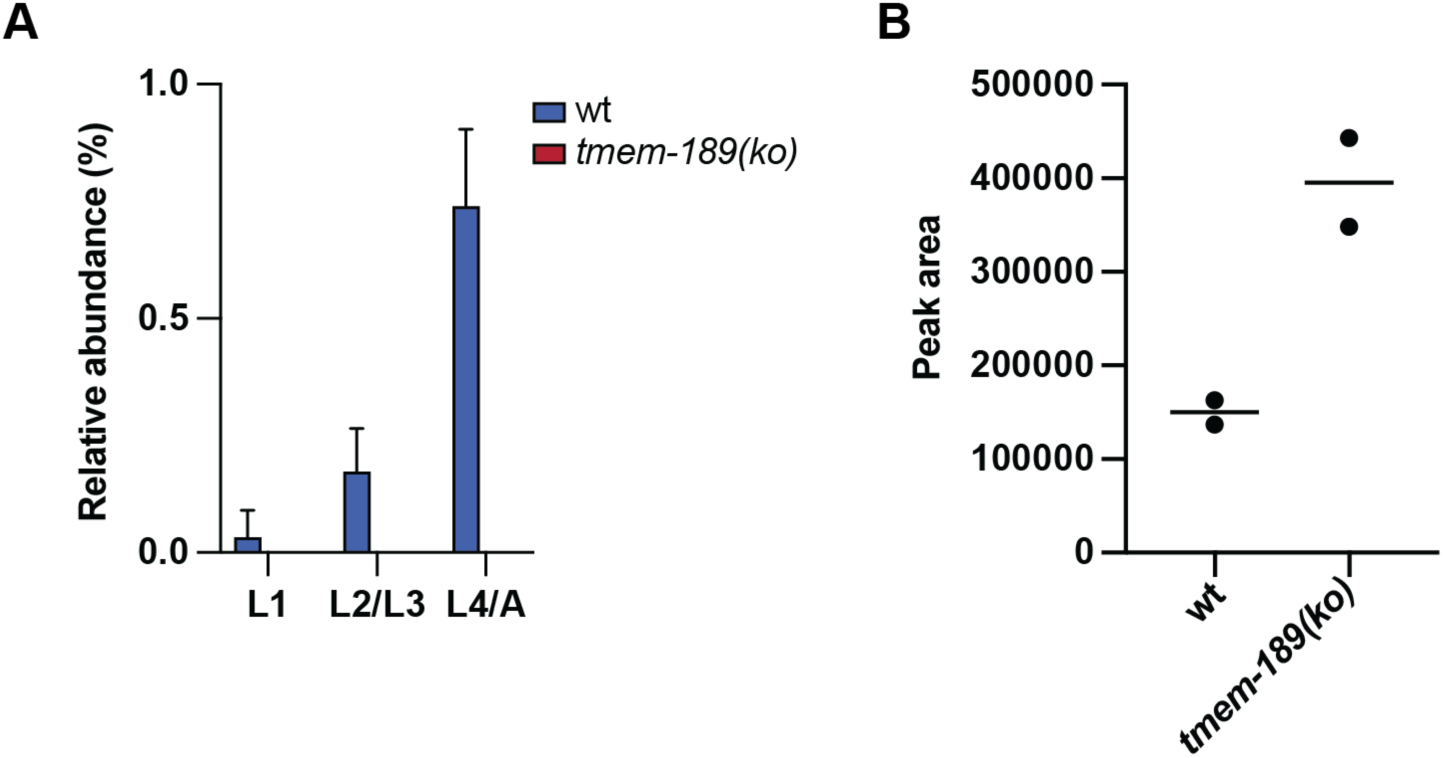
*tmem-189(ko) C. elegans* strain lacks plasmalogens and accumulates alkyl-GPLs. (A) Plasmalogen levels in the wild type (wt) and *tmem-189(ko)* strains across different *C. elegans* developmental stages (L1, L2/L3, and L4/adult). Data correspond to the most abundant plasmalogen-derived DMA (C18:0-DMA) analyzed by GC-MS, and are the mean ± SD relative to total lipid content (n = 3). (B) LC-MS/MS analysis of the most abundant alkyl-PE species (PE *O-*18:0/18:1, m/z = 730.5756) in wt and *tmem-189(ko)* strains at the L4/adult stage.

To assess neuronal function under sublethal stress, we quantified locomotor behavior, a highly sensitive readout of neuronal integrity in *C. elegans*.^46^ At non-lethal FAC concentrations, the *tmem-189(ko)* worms displayed pronounced locomotor defects compared to wt worms (Figures 7C and 7D). Importantly, this impairment was rescued by the ferroptosis inhibitor ferrostatin-1 (Figures 7C and 7D), indicating that the behavioral defect arises from iron-dependent ferroptosis rather than from nonspecific toxicity. These data demonstrate that plasmalogens act as critical *in vivo* modulators of ferroptosis-related damage. These findings extend the anti-ferroptotic role of plasmalogens beyond cultured cells and underscore their importance in maintaining neuronal function.

## DISCUSSION

Ferroptosis is now a major field of research due to its potential anticancer roles and implication in cell death in various diseases.^47^ Although PUFA-containing GPLs are well established as key substrates for ferroptotic lipid peroxidation, the broader structural determinants that make specific GPLs pro-ferroptotic—beyond the presence of polyunsaturated tails—remain largely unresolved. Due to the large number of possible combinations between head groups, tails, and linkages, membrane GPLs are extremely diverse,^48^ and thus establishing the precise relationship between GPL structure and ferroptosis remains a key challenge. Different studies linked distinct lipids to ferroptosis, including PE with adrenic acid (22:4),^49^ PC with two PUFA tails,^50^ or phosphatidylinositol with AA.^51^ In this context, ether lipids have recently emerged as regulators of ferroptosis, yet their structural diversity has contributed to conflicting models about their promoting or protective function and on the molecular basis of their activity.^6–8^ Some studies showed that the depletion of ether lipid biosynthetic enzymes impairs ferroptosis in mammalian cells, while exogenous delivery of polyunsaturated ether lipids restores it.^7,8^ However, Perez et al. demonstrated that *C. elegans* lacking ether lipids are hypersensitive to dihomo-gamma-linolenic acid-induced ferroptosis, while selective plasmalogen deficiency (*tmem-189* mutants) has no sensitizing effect upon this treatment.^6^ Thus, the pro- or anti-ferroptotic nature of ether lipids seems to be context dependent.

Of note, lipid metabolism constitutes a complex network of interconnected reactions, and thus the depletion of some lipid biosynthetic pathways leads to compensatory changes in other lipids. An example is the upregulation of polyunsaturated diacyl-GPLs in ether lipid deficiency (Figures S1G and S1F). Thus, we reasoned that it is critical to compare comprehensive lipidomics data with ferroptosis sensitivity, in order to link correctly lipid structures to ferroptosis. Our data clarified why ether GPLs appear pro-ferroptotic. From all the data we collected, we found that ferroptosis sensitivity scales well with the degree of polyunsaturation in PE. This was the case even for GNPAT and AGPS deficient cells, which raised the possibility that the ether linkages themselves do not affect ferroptosis per se. Rather, the ether linkages seem to dictate the metabolic bias of GPLs toward a combination of PE head group and PUFA, likely due to substrate preferences of SELENOI and TMEM164. This metabolic bias accumulates pro-ferroptotic GPL structures, which would be unfavorable for cell fitness. This is probably why PEDS1 introduces the anti-ferroptotic vinyl ether linkage as a measure to counterbalance the pro-ferroptotic nature of ether lipid metabolic bias. This raises the question: why does this metabolic bias exist? It was recently shown that the plasma membrane is not only asymmetric in GPL class distribution, but also in the degree of unsaturation, leading to a more fluid inner leaflet.^28^ While the physiological relevance of this fluid inner leaflet has to be established, it is possible that the combination of PUFA and PE head group in ether lipids contributes to this asymmetry of fluidity. Thus, our study raises the possibility that one biological function of ether linkage is to dictate a metabolic bias that would accumulate lipids that fluidify the inner leaflet of the plasma membrane, which sensitizes cells to ferroptosis as a side effect of this major function. In this context, the role of the vinyl ether linkage would be to alleviate this side effect, allowing cells to have a polyunsaturated inner leaflet without being too ferroptosis-prone. We believe that the apparent discrepancy in the literature regarding ether lipids and ferroptosis is due to their dual function, and that the anti- and pro-ferroptotic aspects of ether lipids depend on the context and experimental conditions.

The protective function of plasmalogens is particularly evident in neurons, where PE and PUFA containing ether lipids accumulate. SELENOI deficiency causes neurodevelopmental disorders in humans,^31^ which shows that ether PE is particularly important in neurons. As SELENOI is a key enzyme for ether lipid metabolic bias, neurons strongly rely on PEDS1 to prevent the pro-ferroptotic build-up of polyunsaturated ether PE, which could be part of the mechanism how PEDS1 deficiency causes neurodevelopmental dysfunctions.^30^ Indeed, we showed that plasmalogen loss compromises neuronal survival under physiological mild oxidative stress. Along similar lines, plasmalogen deficiency sensitizes *C. elegans* to iron-dependent locomotion impairment, which can be rescued by ferrostatin, a phenotype likely related to neuronal damage.

Our study on neuronal ferroptosis led to the unexpected finding linking plasmalogens and mitochondrial functions. We found that plasmalogen depletion disturbs mitochondrial respiration, increasing ROS production and susceptibility to ferroptosis in neurons. These phenotypes arise independently from ferroptosis induction, indicating that plasmalogen loss alone is sufficient to compromise mitochondrial homeostasis. While the protective role of the vinyl-ether bond in scavenging ROS is well recognized,^11^ the impaired mitochondrial functions observed upon plasmalogen loss points to an additional function in sustaining mitochondrial efficiency and structural integrity. Recent structural studies using electron tomography have shown that cristae adopt a range of morphologies, including amorphous and irregular, non-lamellar forms, which are commonly associated with altered mitochondrial function.^52^ We observed a similarly irregular cristae organization in PEDS1 mutant neurons, which could contribute to the other mitochondrial defects we found. Others showed that in pancreatic ductal adenocarcinoma, monounsaturated ether GPLs localize to mitochondria, where they stabilize respiratory supercomplexes and suppress lipid peroxidation.^8^ Ablating ether lipid synthesis sensitizes these cells to oxidative damage—a defect rescued specifically by MUFA-ether lipids, but not free MUFAs.^8^ Thus, ether lipids have important roles in mitochondria. While proteins embedded in the inner mitochondrial membrane are considered major drivers of cristae shape, membrane lipids play critical roles too. For example, the lipid with a negative spontaneous curvature cardiolipin is required to maintain proper cristae architecture.^53^ Recently, plasmalogen PE has been shown to have a strong negative spontaneous curvature, which was stronger than the effect of diacyl-PE.^54^ Thus, we speculate that plasmalogen PE could be an additional membrane-shaping lipid in the inner mitochondrial membrane. However, further work will be required to determine the mechanisms by which changes in lipid composition give rise to the structural alterations seen in PEDS1 deficiency.

Here we propose that plasmalogens have dual roles: they regulate mitochondrial function and scavenge reactive oxygen species, possibly in the plasma membrane. In their absence, mitochondrial performance is compromised, leading to increased ROS production. At the same time, without the ROS-scavenging capacity of plasmalogens, this oxidative burden overwhelms the cell’s antioxidant defenses. It is therefore plausible that their protective effect in ferroptosis reflects both direct chemical reactivity and indirect regulation of mitochondrial homeostasis. Taken together, these findings refine our understanding of the role of ether lipids in ferroptosis. Rather than acting as a uniform lipid class, ether phospholipids exhibit divergent, context-dependent functions shaped by their bond type, fatty acid composition, and subcellular localization.

## MATERIALS AND METHODS

### Statistics

Statistical tests were performed using GraphPad Prism, R, or Python. Details of the tests and results are provided in supplementary spreadsheet Data S1.

### Cell culture

HeLa cells (HeLa MZ clones) were kindly provided by Jean Gruenberg, University of Geneva. Cells were cultured in Dulbecco’s Modified Eagle Medium (high glucose DMEM, GlutaMAX, pyruvate, Gibco) with 10% fetal calf serum (Eurobio Scientific) and 1% ZellShield cell culture contamination preventive (Minerva-Biolabs) at 37°C and 5% CO_2_. Passages of cells were performed at 80-90% confluency and in the range of 1:5 to 1:20. Cells were washed with Dulbecco’s Phosphate-Buffered Saline (DPBS without calcium/magnesium, Gibco) two times and detached with trypsin-EDTA (Gibco), which was stopped by the addition of complete medium. Cells were counted using Countess II automated Cell Counter (Invitrogen) following manufacturer instructions.

Human iPSCs expressing doxycycline-inducible NGN2 integrated at the AAVS1 safe-harbor locus^55^ (UCSFi001-A; kindly provided by Michael E. Ward, NIH/NINDS) were cultured on Matrigel (Corning® Matrigel® hESC-Qualified Matrix, Product Number 354277) coated plates in mTeSR Plus with supplement at 37 °C and 5% CO₂. Neuronal induction followed the study of Fernandopulle et al.^55^ Cells were dissociated with Accutase and plated on Matrigel coated plates in DMEM/F12 containing N2 supplement, non-essential amino acids, doxycycline (2 µg/mL), and ROCK inhibitor (Y-27632, first 24 h only). After 3 days, cells were replated onto poly-D-lysine (PDL)–coated plates and matured in BrainPhys medium with NeuroCult™ SM1 Neuronal Supplement supplemented with BDNF (10 ng/mL), NT-3 (10 ng/mL), laminin, and doxycycline (+AO). Medium was replaced every 3–4 days. Mild oxidative stress was induced by culturing neurons in antioxidant-free NeuroCult™ SM1 Neuronal Supplement (–AO; catalog #05732, STEMCELL Technologies).

### Lipidomics

HeLa cells were seeded at 750,000 cells on 6 cm dishes and cultured for 24 h. Cells were then washed with ice-cold PBS and scraped in 500 µL of ice-cold PBS, pelleted by centrifugation at 2,500 rpm at 4°C, and snap-frozen in liquid nitrogen. iPSCs were plated in Matrigel-coated 6-well plates and grown to ∼80% confluency. iNs were plated on PDL-coated 6-well plates and cultured at 1.5 million cells per well until day-5 or day-14. All plates were washed with PBS and directly frozen at −80 °C until processing.

HeLa cell pellets were resuspended in 200 µL water followed by the addition of 500 µL methanol. iPSCs and iN were scraped in 100 µL water + 360 µL methanol; an additional 100 µL water and 140 µL methanol were added to iPSC and iN suspensions. Internal standards (4 µL SPLASH LIPIDOMIX Mass Spec Standard plus 20 pmol each of C17 ceramide, C17 Glucosylceramide, and C17 Lactosylceramide, or 4 µL SPLASH LIPIDOMIX Mass Spec Standard plus 2 µL SphingoSPLASH I, all from Avanti Polar Lipids) were added, followed by 250 µL of chloroform. The mixture was shaken for 10 min. 250 µL of water and 250 µL of chloroform were added to induce phase separation and the mixture was shaken for 10 min. Samples were centrifuged for 15 min at 3,000 rpm, and 400 µL of organic phase was transferred to new glass tubes and dried under nitrogen. Finally, samples were reconstituted in 60 µL isopropanol : methanol : water (5:3:2). Three µL of extracts were injected into an Ultimate 3000 UHPLC system coupled to a Q Exactive mass spectrometer (Thermo Fisher Scientific). Lipids were separated with an Accucore C18 column (150 x 2.1 mm, 2.6 µm particles), using as mobile phase solvent A (acetonitrile : water (1:1, v/v) supplemented with 10 mM ammonium formate and 0.1% formate) and solvent B (isopropanol : acetonitrile : water (88:10:2, v/v) supplemented with 2 mM ammonium formate and 0.02% formate) at a 400 µL/min flow rate. The gradient was linear as follows: 35% solvent B at 0 min, 60% B at 4 min, 70% B at 8 min, 85% B at 16 min, 97% B at 25 min, and 100% B at 25.1 min until 31 min. The column was reconditioned at 35% B for 4 min. Data was acquired by data dependent MS2 in positive and negative ion modes through separate injections. MS1 was acquired at 70,000 resolution (at m/z = 200) to cover the range of m/z = 250-1200 with an AGC target of 1,000,000 and 250 msec maximum injection time. Fifteen precursor ions were selected for MS2 analysis in an isolation window of m/z = 1 and fragmented by higher-energy collisional dissociation with a normalized collision energy of 25% and 30% for positive mode and 20%, 30%, and 40% for negative mode. For MS2 analysis, AGC target was 100,000 and maximum injection time was 80 msec, with a resolution of 35,000 at m/z = 200.

Peaks were annotated with MS-DIAL5^56^ based on MS2 spectra, which were further curated based on retention times. Discrimination of alkyl- and alkenyl-GPLs was based on retention time, with alkenyl-GPLs eluting 0.2-0.3 min earlier than the corresponding alkyl-GPLs. Lipids semi-quantification was done by dividing peak areas with those of internal standards and multiplying with the quantities (in pmol) added to samples.

Lipidomics data are provided in supplementary spreadsheet Data S2-S8.

### Mutant cell generation

HeLa cells were mutated using the CRISPR-Cas9-based GENF strategy as previously described.^57^ Briefly, guide RNAs (gRNAs) against the target genes were selected based on high specificity and efficiency using UCSC genome browser. Selected gRNAs were cloned into pX330 plasmid encoding the Cas9 endonuclease. To generate the mutant cells, a plasmid containing the specific gRNA was co-transfected with a plasmid containing HPRT1 gRNA (in a 100:1 ratio) into HeLa MZ cells using lipofectamine 3000 (Thermo Fisher). Five days post-transfection, cells were cultured in media containing 6-thioguanine (6TG, 6 µg/mL) for an additional seven days, allowing the selection of HPRT1 mutated cells (thus cells with high CRISPR-Cas9 efficiency). After the selection, genomic DNA was isolated from the surviving cells and gene-specific primers were used to amplify the region encompassing the Cas9 cleavage site. The efficiency of knockdown was calculated using the Tracking of Indels by DEcomposition tool (TIDE).^58^

List of gRNAs used for GENF strategy:

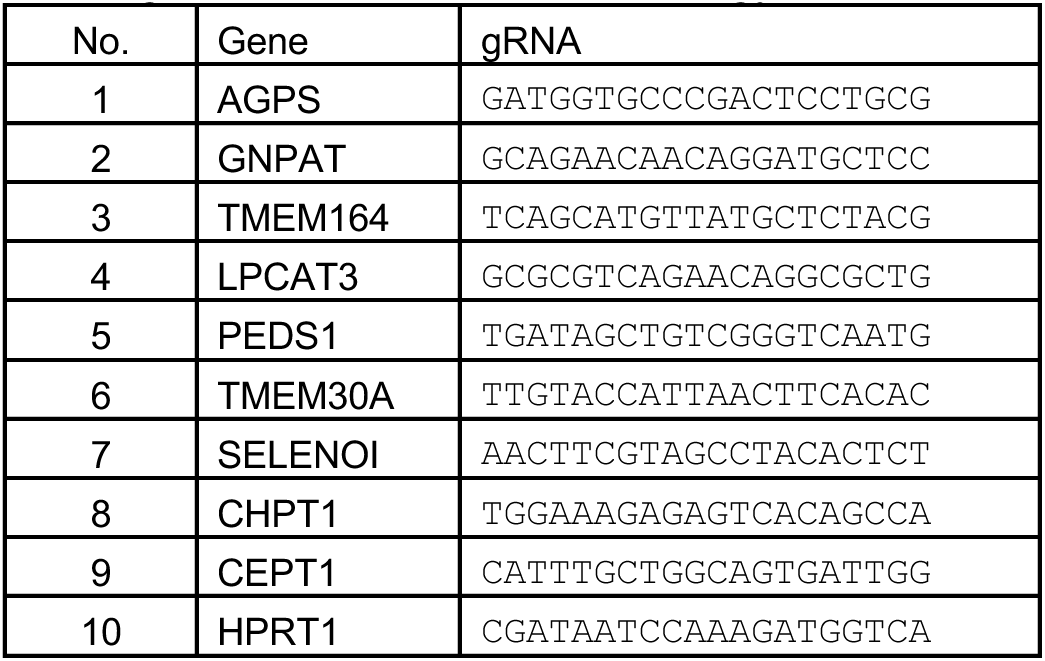

PEDS1 was mutated in human iPSCs cells by CRISPR-Cas9, using pLentiCRISPRv2 plasmid coding for Cas9 and the sgRNA targeting CCTGCTGCTGGCCCGCTGGG (synthesized by GenScript). iPSCs were transfected with Lipofectamine 3000 and selected with puromycin (1 µg/mL). Genomic DNA was extracted by SDS–proteinase K lysis, amplified (1081 bp amplicon), and purified with ExoSAP-IT. Sanger sequencing was analyzed using TIDE analysis to quantify indels, and clones containing frameshift mutations were expanded.

### iPSC Genetic Analysis

Human iPSCs, including control lines and PEDS1 mutant lines, were cultured in 6-well plates and harvested at approximately 70–80% confluency. For each analysis, half of a 6-well plate per cell line was used for genomic DNA extraction. Genomic DNA was isolated using the Wizard® SV Genomic DNA Purification System (Promega) according to the manufacturer’s instructions. DNA concentration and purity were assessed using a NanoDrop™ spectrophotometer.

Genetic stability was evaluated using the hPSC Genetic Analysis Kit (STEMCELL Technologies), a quantitative PCR-based assay designed to detect recurrent chromosomal abnormalities in human pluripotent stem cells. The kit includes primer–probe mixes targeting chromosomal regions 1q, 4p, 8q, 10p, 12p, 17q, 18q, 20q, and Xp. A female diploid genomic DNA control supplied with the kit was used as the reference. For each reaction, 300 ng of genomic DNA was analyzed, and all procedures were performed according to the manufacturer’s protocol.

Ct values were exported and analyzed using the comparative ΔΔCt method. For each sample, ΔCt values were calculated by subtracting the average Ct of the Chr 4p (control chromosome) assay from the Ct of each target locus. ΔΔCt values were then obtained by subtracting the average ΔCt of the genomic DNA control from the ΔCt of each test sample.

Copy number was calculated as:

Copy number = (2−ΔΔCt)×2

Copy number values < 1.8 or > 2.4 with p < 0.05 were considered indicative of potential chromosomal abnormalities. Statistical significance was assessed using one-way ANOVA with Tukey post-hoc testing or unpaired t-tests, as appropriate. The data are provided in supplementary spreadsheet Data S9.

### Ferroptosis assay

To evaluate ferroptosis sensitivity, cells were seeded at a density of 5,000 cells per 100 µL per well in 96-well plates and incubated for 24 h at 37°C in a humidified atmosphere containing 5% CO₂. Cells were then treated with serial dilutions of ML210, prepared in DMSO, to reach final concentrations of 0.001, 0.01, 0.02, 0.04, 0.06, 0.08, 0.1, 0.25, 0.5, 1, 2.5, 5, and 10 µM. Each concentration was tested in quadruplicate. The final concentration of DMSO was maintained at 0.1% (v/v) in all wells, including the control samples. Cells were incubated with ML210 for 24 h. Following treatment, the medium was carefully aspirated and replaced with 100 µL of MTT solution (0.5 mg/mL in culture medium). Plates were incubated for 3 h under standard culture conditions to allow for MTT reduction by metabolically active cells. Subsequently, the MTT-containing medium was removed, and 100 µL of DMSO was added to each well to solubilize the resulting formazan crystals. Absorbance was measured at 595 nm using a Multiskan FC Microplate Photometer (Thermo Scientific). Relative cell viability was calculated by normalizing absorbance values to the DMSO control group. Data analysis and curve fitting were performed using GraphPad Prism software.

### Plasmid construction and stable cell selection

The empty plasmid backbone PB-CAG-IRES-Puro for gene expression in mammalian cells was generated as follows. The IRES-PuroR fragment was amplified from pCAGGS-flpE-puro (Addgene #20733 deposited by Rudolf Jaenisch) by PCR using PrimeSTAR Max with the primers TAATTGCTAGCTGATCTAGAGTCGAGTTAAT and TAAATGGATCCCTGATCAGCGAGCTCT-`AGAT`, thus appending NheI and BamHI restriction sites on the amplicon. The amplicon was incorporated into the corresponding sites of PB CAG-sfGFP (Addgene #133568 deposited by Mario Capecchi), thus replacing the sfGFP-WPRE insert with IRES-PuroR between the CAG promoter and the bGH poly(A) signal. As a consequence, we obtained a plasmid allowing the expression of genes from the CAG promoter, which is amenable for stable integration into the genome with the PiggyBac transposon system, and selection by puromycin. We then inserted between the NheI and EcoRI sites of this backbone HA tagged human codon-optimized human TMEM164 or Saccharomyces cerevisiae Cds1 (ScCds1) made by gene synthesis (GeneArt strings from Thermo Fisher). To generate a plasmid for endoplasmic reticulum labeling, we inserted an mNeonGreen-Sec61b fragment between the NheI and EcoRI sites of PB-CAG-IRES-Puro. The insert was generated by overlap extension PCR with the primers GTCCAGCTAGCCACCATGGTGAGCAAGGGCGAGGA and ACCAGGCATCTTGTACAGCTCGTCCATG-`CC` to amplify mNeonGreen with an NheI site at the 5’ region and a fragment of Sec61b at the 3’ region, and GAGCTGTACAAGATGCCTGGTCCGACCCCCA and GACGGTGAATTCCTACGAACGA-GTGTACTTGCCC to amplify Sec61b with a fragment of mNeonGreen at the 5’ region and an EcoRI site at the 3’ region. The two fragments were assembled through a second round of PCR using the external primers, and inserted between the NheI and EcoRI sites PB-CAG-IRES-Puro. ScCds1 stable cells were generated by co-transfecting HeLa cells with the aforementioned expression plasmid and pCMV-hyPBase^59^ (kindly provided by the Wellcome Sanger institute), which allowed the expression of a transposase that stably integrates the transgene (in this case ScCds1-IRES-PuroR) into the genome. Stable cells were selected with puromycin (0.5 µg/mL) for 14 days.

### Fluorescence imaging and flow cytometry

For TMEM164 localization analysis, HeLa cells were transfected transiently with lipofectamine 3000 to express mNeonGreen-Sec61 and HA-hTMEM164 from the above plasmids. Two days after transfection, cells were fixed with 4% paraformaldehyde in PBS for 30 min. Cells were then washed thrice with PBS and permeabilized with 0.1% Triton X-100 in PBS for 10 min. Cells were washed again in PBS, followed by incubation with primary antibody against HA tag (1:1000 dilution, Enzo cat.# ENZ-ABS118) in PBS containing 1% BSA for 1 h. Cells were washed again with PBS and incubated with an Alexa 594-tagged secondary antibody for 30 min in PBS with 1% BSA. After the incubation, cells were washed and stained with DAPI (200 ng/mL) for 10 min in PBS. Cells were washed again and imaged by confocal microscopy (Leica SP8).

For the analysis of cell surface phosphatidylserine exposure, HeLa cells (control and TMEM30A mutants) were grown in 6 cm dishes for two days. Cells were detached and washed with PBS. One million cells were resuspended in 100 µL of Annexin-binding buffer and mixed with 2.5 µL of APC-Annexin V (BD cat. # 550474) and incubated at room temperature for 15 min. After incubation, 200 µL of binding buffer containing 2.5 µL of propidium iodide (Sigma-Aldrich cat. # P4864) was added, and the sample was analyzed by flow cytometry (BD LSRII Fortessa).

To analyzed membrane fluidity, cells were incubated with 40 nM NR12S^27^ probe for 15 min at 37^°^C. Images were acquired using a laser scanning confocal microscope (Carl Zeiss LSM 780) with an excitation laser at 488 nm, through a 63x/1.4 oil immersion objective. Fluorescence emission images were taken from 550-600 nm (ordered channel, OC) and 600-650 nm (disordered region, DC). Generalized polarization (GP) values were obtained from 32 bits images calculated according to the formula: GP = (OC - DC) / (OC + DC), using an in-house Fiji/ImageJ macro program.

For imaging of iN cells, day-3 neurons were seeded into poly-D-lysine (PDL)–coated ibidi 96-well plates for high-content fluorescence imaging and maintained until maturation. For flow-cytometry experiments, neurons were seeded in PDL-coated 6-well plates.

For live-cell imaging, cells were incubated with fluorescent probes for 60 min at 37 °C. The following dyes were used: 20 nM TMRE to assess mitochondrial membrane potential and 500 nM MitoSOX Red to detect mitochondrial superoxide. All imaging assays were performed in parallel either in standard antioxidant-containing medium (+AO) or in antioxidant-free medium (–AO), as indicated.

For neuronal survival assays based on MAP2 staining, neurons were fixed with 4% paraformaldehyde (PFA) and permeabilized with PBS containing 1% bovine serum albumin (BSA) and 0.05% saponin. Cells were incubated with anti-MAP2 antibody (clone AP20, MAB3418-25UG, Sigma-Aldrich Chemie GmbH). Staining and treatments were performed in +AO or –AO medium. Where indicated, rescue experiments were carried out using 10 µM Ferrostatin-1, 10 µM Z-VAD-FMK, or 100 nM MitoQ.

High-content images were acquired using an Operetta High-Content Imaging System (PerkinElmer). Image quantification—including fluorescence-intensity measurements, cell segmentation, and frequency-distribution analyses—was performed using QuPath software.

Lipid peroxidation imaging using C11-BODIPY 581/591 was performed on a Leica SP8 confocal microscope together with MitoTracker Deep Red to visualize mitochondrial localization of lipid peroxidation under +AO or –AO conditions.

For flow cytometry, lipid peroxidation and cellular reactive oxygen species (ROS) were assessed by staining cells with 5 µM Liperfluo or 5 µM CellROX Deep Red for 30 min at 37 °C. Cells were detached using Accutase, passed through a 40 µm cell strainer, and then analyzed on a BD LSRFortessa flow cytometer. Flow-cytometry data analysis, including gating strategies and population statistics, was performed using FlowJo and Kaluza software.

### RNA-sequencing

Total RNA was isolated from one well of a 6-well plate of mature human iPSC-derived neurons using the RNeasy Kit (QIAGEN), according to the manufacturer’s instructions. RNA concentration was measured using a Qubit fluorometer (Thermo Fisher Scientific), and RNA integrity was assessed using a Bioanalyzer (Agilent Technologies). For library preparation, 500 ng of total RNA was used as input for the TruSeq Stranded mRNA Library Preparation Kit (Illumina). Library molarity and quality were assessed using a Qubit fluorometer and a TapeStation system with a DNA High Sensitivity chip (Agilent Technologies). Libraries were loaded for clustering on a single-read Illumina flow cell, and sequencing was performed on an Illumina HiSeq 4000 platform using TruSeq SBS chemistry to generate single-end reads of 100 bp. RNA sequencing was carried out at the iGE3 Genomics Platform of the University of Geneva.

Sequencing reads in FASTQ format were aligned to the human reference genome (GRCh38) using STAR (v2.7.10b).^60^ Gene-level read counts were generated from the resulting BAM files using HTSeq-count,^61^ with genes defined according to the Ensembl v108 annotation and using the reverse-strand option. Genes with fewer than 10 raw counts in at least three samples were excluded from downstream analysis, resulting in 16,625 expressed genes retained. Differential gene expression analysis was performed using DESeq2^62^ (v1.36.0) with default parameters. Genes were considered differentially expressed if they exhibited an absolute log2 fold change ≥ 1 (corresponding to a ≥ 2-fold change) and a false discovery rate (FDR) ≤ 0.05. Bioinformatic analyses were performed with support from the Bioinformatics Competence Center (UNIL-EPFL). The results are provided in supplementary spreadsheet Data S10.

### qPCR analysis

Total RNA was isolated using the RNeasy Mini Kit. cDNA synthesis was performed using the iScript cDNA kit (Bio-Rad). qPCR reactions were set up using PowerUp SYBR Green Master Mix and run on a QuantStudio 6 system. Expression was normalized to GAPDH using the ΔΔCt method. Primer sequences used are listed below:

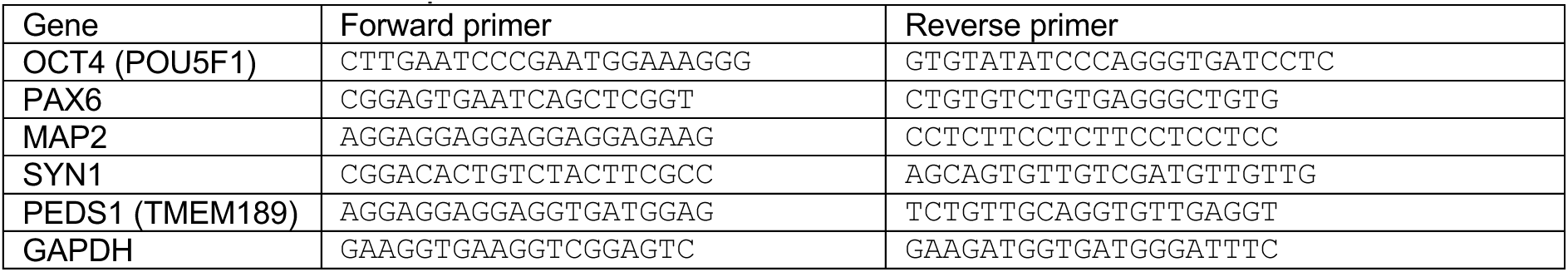

### Crude mitochondrial fraction isolation

Crude mitochondrial fractions were isolated from mature NGN2-induced neurons. Briefly, ∼7 × 10⁶ neurons were washed in ice-cold PBS and scraped into mitochondrial isolation buffer (10 mM Tris-HCl pH 7.4, 250 mM sucrose, 1 mM EDTA, protease inhibitors). Cells were homogenized on ice by 30 passes through a 25-gauge needle. Nuclei and debris were removed by sequential centrifugation at 300 × *g* for 5 min and 600 × *g* for 5 min at 4°C. The resulting supernatant was centrifuged at 13,000 × *g* for 10 min to pellet crude mitochondria. Pellets were washed once in isolation buffer and processed immediately for Blue Native PAGE or proteomic analysis.

### Proteomics sample preparation for mass spectrometry

Mass spectrometry-based proteomics-related experiments were performed by the Proteomics Core Facility at EPFL. Each sample was digested by filter aided sample preparation (FASP)^63^ with minor modifications.

Proteins (25 ug) were reduced with 10 mM TCEP in 8M Urea, 0.1M Tris-HCl pH 8.0 at 37°C for 60min and further alkylated in 40mM iodoacetamide at 37°C for 45min in the dark. Proteins were digested overnight at 37°C using 1/50 w/w enzyme-to-protein ratio of mass spectrometry grade Trypsin Gold and LysC. Generated peptides were desalted in StageTips using 6 disks from an Empore C18 (3 M) filter based on the standard protocol.^64^ Purified peptides were dried down by vacuum centrifugation.

### LC-MS/MS for proteomics

Samples were resuspended in 2% Acetonitrile, 0.1% Formic acid and nano-flow separations were performed on a Vanquish Neo nano UPLC system (Thermo Fischer Scientific) on-line connected with an Exploris 480 Orbitrap Mass Spectrometer. A capillary precolumn (Acclaim Pepmap C18; 3 μm-100 Å; 2 cm x 75 μm ID) was used for sample trapping and cleaning. Analytical separations were performed at 250 nL/min over a 150 min biphasic gradients on a 50 cm long in-house packed capillary column (75 μm ID; ReproSil-Pur C18-AQ 1.9 μm silica beads; Dr. Maisch).

Mass spectrometry data were acquired on an Orbitrap Exploris mass spectrometer (Thermo Fisher Scientific) operated in data-independent acquisition (DIA) mode. Full MS1 scans were acquired in the Orbitrap over an m/z range of 420-680 at a resolution of 60,000 (at m/z 200), with a normalized automatic gain control (AGC) target of 300% and a maximum injection time set at auto. MS1 scans were followed by sequential DIA MS2 scans covering an m/z range of 430-670, using non-overlapping 6.66 m/z isolation windows resulting in 36 DIA windows per cycle. Fragmentation was performed by higher-energy collisional dissociation (HCD) using a normalized stepped collision energy of 22, 26, 30. MS2 spectra were acquired in the Orbitrap at a resolution of 45,000 (at m/z 200), with a normalized AGC target % at 3000.

### Data Analysis for proteomics

Raw data files were analyzed with DIA-NN^65^ (version 1.9.2) using library free search against Swissprot_Human_20418Sequences_LR2024_01 database, complemented with the common MaxQuant contaminants.^66^ Results were filtered at 1% FDR. Search parameters included: minimum fragment m/z at 200, maximum fragment m/z at 1800, N-terminal methionine excision enabled, in silico digest with cuts at K* and R*, maximum number of missed cleavages set to 1, minimum peptide length set to 7, maximum peptide length set to 30, minimum precursor m/z at 300, maximum precursor m/z at 1800, minimum precursor charge set to 1, maximum precursor charge set to 4. Carbamidomethylation (C) was set as a fixed modification, whereas oxidation (M) as a variable modification. Maximum number of variable modifications was set to 2 and Match between runs (MBR) was enabled. The Unrelated runs option was checked. Resulting DIANN report was converted to MSstats format using MSstatsConvert (v1.18.1) and analyzed in R (v.4.5.1) with MSstats (v4.16.1).^67^ Differential protein expression analysis was performed using the linear mixed-effects model followed by the Benjamini–Hochberg procedure. The data are provided in supplementary spreadsheet Data S11.

### Mitochondrial respiration

Mitochondrial respiration was measured using the Agilent Seahorse XF Mito Stress Test Kit (103015-100, Bucher Biotec AG) on an Agilent Seahorse XFe96 Analyzer, following the manufacturer’s instructions. HeLa cells (10,000 cells/well) or day-3 neurons (30,000 cells/well) were plated on PDL-coated XF96 plates. On the day of the assay, cells were incubated for 60 min in Seahorse DMEM (10 mM glucose, 2 mM glutamine, 1 mM pyruvate, pH 7.4) in a non-CO₂ incubator. The sequential injections were: Oligomycin (1 µM), FCCP (1 µM for HeLa, 3 µM for neurons) and Rotenone/Antimycin A (0.5 µM each). OCR values were normalized to total protein measured by BCA assay.

### ATP measurement

Cellular ATP levels were quantified using the CellTiter-Glo Luminescent Cell Viability Assay (Promega). Day-3 neurons were plated on poly-D-lysine–coated black 96-well plates and maintained under the indicated experimental conditions. At the time of measurement, an equal volume of CellTiter-Glo reagent was added directly to each well containing cells and culture medium. Plates were incubated for 10 min at room temperature with gentle shaking to allow complete cell lysis and stabilization of the luminescent signal. The lysates were then transferred to white 96-well plates and luminescence was measured using a microplate reader.

### Blue Native PAGE

For Blue Native PAGE, mitochondrial pellets were solubilized in digitonin-containing NativePAGE lysis buffer (Thermo Fisher Scientific) on ice for 10–15 min and clarified by centrifugation at 20,000 × g for 30 min at 4 °C. The supernatant was supplemented with NativePAGE G-250 sample additive (¼ of detergent concentration) and resolved on NativePAGE Novex 4–16% Bis-Tris gels (Thermo Fisher Scientific).

Electrophoresis was performed at 150 V, initially using dark cathode buffer (containing Coomassie G-250), followed by light cathode buffer (reduced G-250 concentration) after dye-front migration to improve protein resolution. Proteins were transferred to PVDF membranes and immunoblotted for respiratory chain complex subunits using antibodies against NDUFB8 (Complex I) and UQCRC1 (Complex III). Detection was performed using standard chemiluminescence methods.

### Transmission Electron Microscopy (TEM) and Electron Tomography

Day-14 neurons, including control lines and PEDS1 mutant lines, were cultured in glass bottomed Petri dishes and fixed overnight for electron microscopy in a solution of 1.2% glutaraldehyde and 2% paraformaldehyde in 0.1 M phosphate buffer, at pH 7.4. They were then post-fixed in 1.5% potassium ferrocyanide and 2% osmium, followed by 1% thiocarbohydrazide and then 2% osmium tetroxide alone, and then overnight in 1% uranyl acetate. The next day they were washed in distilled water at 50°C, before being stained again with lead aspartate at the same temperature. They were finally dehydrated through increasing concentrations of alcohol and then embedded in Durcupan ACM (Fluka, Switzerland) resin. The Petri dishes were then filled with a 3 mm deep layer of pure resin, and hardened for 24 h in a 65°C oven. For preparing sections for transmission electron microscopy, the glass bottom of the dish was released from the rest of the resin by immersion in liquid nitrogen revealing the resin surface and the embedded cultured cells. Using a light microscopy, regions of resin containing cells of interest were cut away from the rest and mounted on blank resin blocks with acrylic glue. The final block was trimmed with glass knives to form a face ready for serial sectioning. Short series of around 20 sections (50 nm thick) were cut with a diamond knife mounted in an ultramicrotome (Leica UC7), and collected onto single-slot, copper grids with a pioloform support film. These sections were contrasted with lead citrate and uranyl acetate and images taken using an FEI Spirit TEM with Eagle CCD camera.

Electron tomography was performed on mitochondria of interest on the 50 nm resin sections collected on the same single slot grids. Tilt series were acquired using a FEI Tecnai F20 transmission electron microscope operated at 200 kV. Images were recorded over an angular range of −70° to +70° using 2° increments. Tilt series alignment and tomographic reconstruction were carried out using standard weighted back-projection. The movies of reconstituted tomographs are provided in Data S12 and S13.

### *C. elegans* strains and culture methods

We followed standard procedures for growing and maintaining *C. elegans* strains on Nematode Growth Medium (NGM), with *Escherichia coli* OP50 as food source.^68^ The nematode-rearing temperature was set at 20^°^C for all of our experiments and nematode populations were synchronized using hypochlorite bleaching of gravid adults following standard protocol.^69^ The following strains were used: the N2:wild-type (WT) hermaphrodite Bristol isolate was purchased from Caenorhabditis Genetics Center (CGC) and *tmem-189(ko)* was generated in this work as described below.

### Generation of the *C. elegans tmem-189(ko)* mutant

The *tmem-189*/Y53C10A.5*(cer184)* deletion mutant strain, also referred to in this manuscript as *tmem-189(ko)*, was generated by CRISPR/Cas9-mediated genome editing in an attempt to create an endogenous reporter by Nested CRISPR, following a protocol previously described.^70^ Briefly, two 20-nucleotide CRISPR RNAs (crRNAs) targeting the first and last exons of the gene (5’-CTTCAGAATGACATCCAGTT-3’ and 5’-GGCGGGGGGTTACTGTAACT-3’), along with the *dpy-10* co-injection marker, trans-activating CRISPR RNA (tracrRNA) and SpCas9, were injected into young adult hermaphrodites (P_0_), which were then individually separated for recovery and to lay F_1_ progeny. F_1_ worms with the dumpy phenotype were then separated and allowed to lay F_2_ progeny. These F_1_ worms were then genotyped through single-worm PCR using primers that target the flanking regions of the deleted sequence. Eight worms from the F_2_ progeny of candidate F_1_ worms carrying the deletion were individually transferred onto NGM plates to isolate homozygous individuals. Finally, the edited gene was amplified by PCR and sequenced through Sanger sequencing. We found that *tmem-189(cer184)* is an indel allele with the following sequence (*tmem-189* initiation codon, original *tmem-189* STOP codon now out-of-frame, and downstream in-frame STOP codon are indicated in capitals): ATGacatcattgacattcagaacgacagTAAccccccgcctgccttctcgcctacagtaccccaaatcattgatctcatctcatatgtctacttgcgcTAG, and producing a 32-amino acid truncated peptide (MTSLTFRTTVTPRLPSRLQYPKSLISSHMYLR). The selected *tmem-189(cer184)* mutant allele was outcrossed twice to WT yielding the strain MEA1 used for phenotypic analyses.

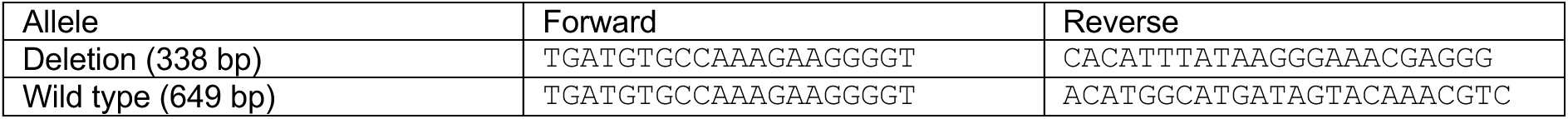

### Lipid analysis in *C. elegans*

To analyze plasmalogens as dimethylacetals (DMAs) in *C. elegans* at various developmental stages, worms were synchronized before collection. L1-worms (∼5,000) were directly collected after egg hatching, and the remaining worms were then transferred to NGM plates seeded with *E. coli* OP50 and incubated at 20 °C for 24 h and 48 h to obtain equivalent numbers of L2/L3 and L4/adult worms, respectively. Worms were collected in M9 buffer, washed twice at 400 x *g* for 2 min to remove *E. coli*, and pellets were stored at -80 °C. The pellets were resuspended in 250 µL of PBS and transferred to Pyrex glass tubes. Samples were then mixed with 8 mL of chloroform/methanol (2:1, v/v) by vortexing and incubated on ice for 1 h. Phase separation was induced by adding 2 mL of 120 mM KCl, followed by vortexing for 10 s and centrifugation at 400 × g for 5 min at 4 °C. After removing the upper aqueous/protein phase, the lower phase was passed through a Whatman No. 1 filter paper (Sigma Aldrich) into a new Pyrex glass tube and dried under a N_2_ stream at 35 °C. To obtain the FAMEs, dried samples were subjected to acid hydrolysis at 55 °C overnight in 1 mL toluene and 2 mL of 1% H_2_SO_4_ in methanol. The reaction was quenched with 2 mL of 0.2 M KHCO_3_, mixed gently with 5 mL hexane:diethyl ether (1:1 v/v) containing 0.01% 2,6-di-tert-butyl-4-methylphenol (BHT; Scharlab) to minimize oxidation and centrifuged at 500 x *g* for 2 min at 4 °C. The upper hexane:ether phase was transferred to a separate tube and the lower phase was re-extracted with 5 mL hexane:diethylether (1:1 v/v). Finally, the combined upper phase was dried with N_2_ at 35 °C and resuspended in 400 µL hexane. For GC-MS analysis, 100 µL of the reaction mix was transferred to 1.5 mL SureStop amber vials (Thermo Fisher Scientific). The analysis was performed using an Agilent 7890B system equipped with an HP-5ms (30 m x 0.25 mm, 0.25 µm) column and a 5977B electron impact mass selective detector. Helium was used as the carrier gas at a flow rate of 1 mL/min. A 2 µL sample was injected in splitless mode, and the column temperature was programmed to ramp at a rate of 6 °C /min from 60 °C to 270 °C, with a 10-min hold at 270°C. The inlet, MSD transfer line, ion source, and quadrupole temperatures were set at 250 °C, 280 °C, 230 °C, and 150 °C, respectively. The electron energy was set at 70 eV, and the mass selective detector operated in scan mode within a mass range of 40 to 600 m/z. Data analysis was performed using Agilent MSD ChemStation (v. F.01.00.1903) software, which enabled quantification of the detected species relative to the total lipids. C18:0-DMA was the most abundant species.

To detect PE O-18:0/18:1, the most abundant alkyl-GPL in *C. elegans*, we performed LC-MS/MS analysis. Total lipid extracts from synchronized L4/adult worms were prepared as described above. Dried extracts were resuspended in hexane and filtered through 0.22 µm PVDF syringe filters (Scharlab). LC-MS/MS analysis was performed using an Agilent platform consisting of a 1290 Infinity II UHPLC system coupled to a 6550 iFunnel Q-TOF mass spectrometer equipped with a Jet Stream electrospray ionization (ESI) source and controlled by MassHunter B.05.00 software. Chromatographic separation was achieved using a ZORBAX Eclipse Plus C18 reversed-phase column (100 × 2.1 mm, 1.8 µm particle size). For negative ion mode analysis, mobile phase A consisted of acetonitrile:water (60:40, v/v) with 10 mM ammonium acetate, and mobile phase B consisted of isopropanol:acetonitrile (90:10, v/v), also containing 10 mM ammonium acetate. The gradient elution (expressed as %B) was as follows: 15% (0 min), 30% (0–2 min), 48% (2–2.5 min), 72% (2.5–8.5 min), 99% (8.5–11.5 min), 99% (11.5–12 min), 15% (12–12.1 min), and 15% (12.1–15 min). The flow rate was maintained at 0.6 mL/min, and the column temperature was set to 60 °C. The injection volume was 5 µL. Mass spectrometric acquisition was performed over a mass range of m/z 100–1700. Source parameters were set as follows: drying gas temperature, 200 °C; drying gas flow, 14 L/min; nebulizer pressure, 35 psi; capillary voltage, +3.5 kV; and fragmentor voltage, 175 V. Initial full-scan MS data were acquired, and selected precursor ions were subsequently subjected to targeted MS/MS fragmentation for structural confirmation. The areas under the curve were determined using the MassHunter quantitative analysis tool.

### *C. elegans* iron-overload assay and survival assessment

The iron-overload assay was performed as previously described.^45^ Briefly, synchronized animals at L4 stage were placed on small plates (Sarstedt, Cat#82.1135.500) containing 3mL of NGM supplemented with 10 μM 5-fluoro-2’-deoxyuridine (FUdR, Cat#F0503) to sterilize worms and prevent progeny from hatching. Different concentrations of filter-sterilized FAC (Sigma, Cat#F5879) dissolved in M9 buffer (ranging from 50 mM to 500 mM) was applied to the surface of NGM plates and stored in the dark for 24 h. The next day approximately 40-50 hermaphrodites (L4 stage) were transferred per plate (control (M9)-or FAC-containing plates) and at least three plates for every condition were used. Death events were monitored up to 96 h after the onset of FAC exposure. Three independent experiments were performed.

### Monitoring *C. elegans* locomotion upon iron-overload

Tracking of *C. elegans* upon iron-overload and co-treatment with anti-ferroptotic agent ferrostatin-1 (Fer-1) was performed using the Fiji plugin (Trackmate).^45,71^ Briefly, synchronized animals at L4 stage were transferred to control- or FAC-containing plates (as described above). Given that the anti-ferroptotic agent ferrostatin-1 (Fer-1) (Sigma, Cat# SML0583) is dissolved in DMSO (AppliChem, Cat# A3672, 0250) we used DMSO-containing plates in the same concentration as control. Fer-1 was initially dissolved in DMSO and then diluted in M9 before plating onto the OP50 food source. The final concentration of FAC, Fer-1, and DMSO in plates were 200 mM, 200 μM, and 1%, respectively. Control and treated plates were always stored in the dark for 24 h before transferring animals onto them. The next day approximately 40-50 hermaphrodites (L4 stage) were transferred per plate and at least three plates for every condition were used. Movies were taken approximately 48h after the onset of FAC or Fer-1 exposure with each recording lasting 40 to 50 sec. All movies were subsequently analyzed using Trackmate.

## Supporting information

Data_S1_Statistics

Data_S2_HeLa-lipidomics

Data_S3_AGPS-GNPAT_Lipidomics

Data_S4_TMEM164-LPCAT3_Lipidomics

Data_S5_ScCds1tg_Lipidomics

Data_S6_CHPT1-CEPT1_Lipidomics

Data_S7_PEDS1-SELENOI_Lipidomics

Data_S8_iNeuron_Lipidomics

Data_S9_genetic_analysis_iPSC

Data_S10_RNA_seq

Data_S11_Proteomics_Mitochondria

Data_S12_Control_Tomo_movie

Data_S13_sgPEDS1_Tomo_movie

## ACKNOWLEDGMENTS

We are grateful to André Nadler and Kristin Böhlig (Max-Planck-Institute of Molecular Cell Biology and Genetics) for contributing with unpublished data, Keiken Ri and Sofía Rodriguez-Gallardo (IPMC) for generating plasmids for subcellular localization analysis, Bruno Antonny and his lab members (IPMC) for support and discussion, the PAB-Azur platform (IPMC) for lipidomics, Sophie Abélanet and the MICA platform (IPMC) for assistance in microscopy and flow cytometry, Nathalie Leroudier (IPMC) for Sanger sequencing, Kosuke Yusa (Kyoto University) for advices related to the PiggyBac transposon system, Francesco Palumbo (EPFL) for assistance with flow cytometry, Graham Knott and Jérôme Blanc (EPFL) for electron microscopy support, and Nicolas Chiaruttini (EPFL) for help with microscopy assistance and quantification.

A.A. was supported by the Deutsche Forschungsgemeinschaft (DFG), grant AS 647/1-1, project number 672757. J.C. was supported by grant PID2023-146930NB-I00 from Agencia Estatal de Investigación (AEI)-Spain. H.T. was supported by the Japan Science and Technology Agency (JST) Exploratory Research for Advanced Technology (ERATO) (JPMJER2101), JST FOREST program (JPMJFR230H), JST NBDC (JPMJND2305), and the JSPS KAKENHI (25H01425, and 25H01426). H.T. and T.H. were supported by JST ASPIRE (JPMJAP2505). D.D. and T.H. were supported by the French Government (National Research Agency, ANR) ACYLOMICS project (ANR-25-CE44-1525). M.E.-A. was supported by grants PID2021-123336NB-C21 and PID2024-158644NB-C21 from Agencia Estatal de Investigación (AEI)-Spain (MCIN/AEI/10.13039/ 501100011033) and “ERDF A way of making Europe”. H.R. was supported by the Swiss National Science Foundation (project number 185898 and 184949). T.H. was supported by CNRS/Inserm ATIP-Avenir, ANR through the “Investments for the Future” IDEX UCAJedi ANR-15-IDEX-01, Inserm International Research Project (AtypicoLipid), and Tokyo University of Agriculture and Technology Global Research Innovation Institute (GIR/ARC teams).

## AUTHOR CONTRIBUTIONS

A.A. conceptualization, investigation (iNeuron analyses), resources, funding acquisition, writing (original draft); G.P.J. conceptualization, investigation (ferroptosis in mutant Hela cells), resources, writing (original draft); I.G. investigation (*C. elegans*); S.S. resources, validation; P.P. investigation (lipidomics); J.A.M. validation; S.H. investigation (native PAGE); O.B. investigation (lipidomics); L.F. investigation (lipidomics); E.B.-M. resources, validation; J.V. resources, validation; J.C. resources, validation, funding acquisition; Y.M. software; F.B. software; J.C. investigation (flow cytometry); H.T. software; D.D investigation (lipidomics); M.E.-A. resources, validation, funding acquisition; H.R. conceptualization, supervision, funding acquisition, project administration; G.D’A. conceptualization, supervision, funding acquisition, project administration, writing (original draft); T.H. conceptualization, investigation (lipidomics), supervision, funding acquisition, project administration, writing (original draft). All authors participated in writing (review & editing).

## REFERENCES

1. Dixon, Scott J., Lemberg, Kathryn M., Lamprecht, Michael R., Skouta, R., Zaitsev, Eleina M., Gleason, Caroline E., Patel, Darpan N., Bauer, Andras J., Cantley, Alexandra M., Yang, Wan S., et al. (2012). Ferroptosis: An Iron-Dependent Form of Nonapoptotic Cell Death. Cell 149, 1060–1072. 10.1016/j.cell.2012.03.042

2. Li, Z., Lange, M., Dixon, S.J., and Olzmann, J.A. (2024). Lipid Quality Control and Ferroptosis: From Concept to Mechanism. Annual Review of Biochemistry 93, 499–528. 10.1146/annurev-biochem-052521-033527

3. Moore, S.A. (2001). Polyunsaturated Fatty Acid Synthesis and Release. Journal of Molecular Neuroscience 16, 195–200. 10.1385/jmn:16:2-3:195

4. Lei, P., Walker, T., and Ayton, S. (2025). Neuroferroptosis in health and diseases. Nature Reviews Neuroscience 26, 497–511. 10.1038/s41583-025-00930-5

5. Mohan, S., Alhazmi, H.A., Hassani, R., Khuwaja, G., Maheshkumar, V.P., Aldahish, A., and Chidambaram, K. (2024). Role of ferroptosis pathways in neuroinflammation and neurological disorders: From pathogenesis to treatment. Heliyon 10, e24786. 10.1016/j.heliyon.2024.e24786

6. Perez, M.A., Clostio, A.J., Houston, I.R., Ruiz, J., Magtanong, L., Dixon, S.J., and Watts, J.L. (2022). Ether lipid deficiency disrupts lipid homeostasis leading to ferroptosis sensitivity. PLOS Genetics 18, e1010436. 10.1371/journal.pgen.1010436

7. Zou, Y., Henry, W.S., Ricq, E.L., Graham, E.T., Phadnis, V.V., Maretich, P., Paradkar, S., Boehnke, N., Deik, A.A., Reinhardt, F., et al. (2020). Plasticity of ether lipids promotes ferroptosis susceptibility and evasion. Nature 585, 603–608. 10.1038/s41586-020-2732-8

8. Cui, W., Liu, D., Gu, W., and Chu, B. (2021). Peroxisome-driven ether-linked phospholipids biosynthesis is essential for ferroptosis. Cell Death & Differentiation 28, 2536–2551. 10.1038/s41418-021-00769-0

9. Gallego-García, A., Monera-Girona, A.J., Pajares-Martínez, E., Bastida-Martínez, E., Pérez-Castaño, R., Iniesta, A.A., Fontes, M., Padmanabhan, S., and Elías-Arnanz, M. (2019). A bacterial light response reveals an orphan desaturase for human plasmalogen synthesis. Science 366, 128–132. 10.1126/science.aay1436

10. Werner, E.R., Keller, M.A., Sailer, S., Lackner, K., Koch, J., Hermann, M., Coassin, S., Golderer, G., Werner-Felmayer, G., Zoeller, R.A., et al. (2020). The TMEM189 gene encodes plasmanylethanolamine desaturase which introduces the characteristic vinyl ether double bond into plasmalogens. Proceedings of the National Academy of Sciences 117, 7792–7798. 10.1073/pnas.1917461117

11. Paul, S., Lancaster, G.I., and Meikle, P.J. (2019). Plasmalogens: A potential therapeutic target for neurodegenerative and cardiometabolic disease. Progress in Lipid Research 74, 186–195. 10.1016/j.plipres.2019.04.003

12. Horta Remedios, M., Liang, W., González, L.N., Li, V., Da Ros, V.G., Cohen, D.J., and Zaremberg, V. (2023). Ether lipids and a peroxisomal riddle in sperm. Frontiers in Cell and Developmental Biology 11, 1166232. 10.3389/fcell.2023.1166232

13. Dorninger, F., Forss - Petter, S., and Berger, J. (2017). From peroxisomal disorders to common neurodegenerative diseases – the role of ether phospholipids in the nervous system. FEBS Letters 591, 2761–2788. 10.1002/1873-3468.12788

14. Harayama, T. (2023). Metabolic bias: Lipid structures as determinants of their metabolic fates. Biochimie 215, 34–41. 10.1016/j.biochi.2023.09.019

15. Koch, J., Lackner, K., Wohlfarter, Y., Sailer, S., Zschocke, J., Werner, E.R., Watschinger, K., and Keller, M.A. (2020). Unequivocal Mapping of Molecular Ether Lipid Species by LC–MS/MS in Plasmalogen-Deficient Mice. Analytical Chemistry 92, 11268–11276. 10.1021/acs.analchem.0c01933

16. Eaton, J.K., Furst, L., Ruberto, R.A., Moosmayer, D., Hilpmann, A., Ryan, M.J., Zimmermann, K., Cai, L.L., Niehues, M., Badock, V., et al. (2020). Selective covalent targeting of GPX4 using masked nitrile-oxide electrophiles. Nature Chemical Biology 16, 497–506. 10.1038/s41589-020-0501-5

17. Conrad, M., and Friedmann Angeli, J.P. (2015). Glutathione peroxidase 4 (Gpx4) and ferroptosis: what’s so special about it? Molecular & Cellular Oncology 2, e995047. 10.4161/23723556.2014.995047

18. Colard, O., Breton, M., and Bereziat, G. (1984). Arachidonyl transfer from diacyl phosphatidylcholine to ether phospholipids in rat platelets. Biochemical Journal 222, 657–662. 10.1042/bj2220657

19. Arafeh, R., Shibue, T., Dempster, J.M., Hahn, W.C., and Vazquez, F. (2024). The present and future of the Cancer Dependency Map. Nature Reviews Cancer 25, 59–73. 10.1038/s41568-024-00763-x

20. Reed, A., Ware, T., Li, H., Fernando Bazan, J., and Cravatt, B.F. (2023). TMEM164 is an acyltransferase that forms ferroptotic C20:4 ether phospholipids. Nature Chemical Biology 19, 378–388. 10.1038/s41589-022-01253-7

21. Hashidate-Yoshida, T., Harayama, T., Hishikawa, D., Morimoto, R., Hamano, F., Tokuoka, S.M., Eto, M., Tamura-Nakano, M., Yanobu-Takanashi, R., Mukumoto, Y., et al. (2015). Fatty acid remodeling by LPCAT3 enriches arachidonate in phospholipid membranes and regulates triglyceride transport. eLife 4, e06328. 10.7554/eLife.06328

22. Horibata, Y., Elpeleg, O., Eran, A., Hirabayashi, Y., Savitzki, D., Tal, G., Mandel, H., and Sugimoto, H. (2018). EPT1 (selenoprotein I) is critical for the neural development and maintenance of plasmalogen in humans. Journal of Lipid Research 59, 1015–1026. 10.1194/jlr.P081620

23. He, Y.L., and Qian, H.W. (2025). Cryo-EM structure of human choline-phosphotransferase 1. PDB Entry, 9UET. 10.2210/pdb9uet/pdb

24. Wang, Z., Yang, M., Yang, Y., He, Y., and Qian, H. (2023). Structural basis for catalysis of human choline/ethanolamine phosphotransferase 1. Nature Communications 14, 2529. 10.1038/s41467-023-38290-2

25. Abramson, J., Adler, J., Dunger, J., Evans, R., Green, T., Pritzel, A., Ronneberger, O., Willmore, L., Ballard, A.J., Bambrick, J., et al. (2024). Accurate structure prediction of biomolecular interactions with AlphaFold 3. Nature 630, 493–500. 10.1038/s41586-024-07487-w

26. Nagata, S., Sakuragi, T., and Segawa, K. (2020). Flippase and scramblase for phosphatidylserine exposure. Current Opinion in Immunology 62, 31–38. 10.1016/j.coi.2019.11.009

27. Kucherak, O.A., Oncul, S., Darwich, Z., Yushchenko, D.A., Arntz, Y., Didier, P., Mély, Y., and Klymchenko, A.S. (2010). Switchable Nile Red-Based Probe for Cholesterol and Lipid Order at the Outer Leaflet of Biomembranes. Journal of the American Chemical Society 132, 4907–4916. 10.1021/ja100351w

28. Lorent, J.H., Levental, K.R., Ganesan, L., Rivera-Longsworth, G., Sezgin, E., Doktorova, M., Lyman, E., and Levental, I. (2020). Plasma membranes are asymmetric in lipid unsaturation, packing and protein shape. Nature Chemical Biology 16, 644–652. 10.1038/s41589-020-0529-6

29. Cobley, J.N., Fiorello, M.L., and Bailey, D.M. (2018). 13 reasons why the brain is susceptible to oxidative stress. Redox Biology 15, 490–503. 10.1016/j.redox.2018.01.008

30. Amelan, A., Collins, S.C., Damseh, N.S., Hamada, N., Salim, A., Dvir, E., Monderer-Rothkoff, G., Harel, T., Nagata, K.-i., Yalcin, B., et al. (2026). CRISPR knockout screens reveal genes and pathways essential for neuronal differentiation and implicate PEDS1 in neurodevelopment. Nature Neuroscience 29, 592–603. 10.1038/s41593-025-02165-0

31. Ahmed, M.Y., Al-Khayat, A., Al-Murshedi, F., Al-Futaisi, A., Chioza, B.A., Pedro Fernandez-Murray, J., Self, J.E., Salter, C.G., Harlalka, G.V., Rawlins, L.E., et al. (2017). A mutation of EPT1 (SELENOI)underlies a new disorder of Kennedy pathway phospholipid biosynthesis. Brain 140, aww318. 10.1093/brain/aww318

32. Tian, R., Abarientos, A., Hong, J., Hashemi, S.H., Yan, R., Dräger, N., Leng, K., Nalls, M.A., Singleton, A.B., Xu, K., et al. (2021). Genome-wide CRISPRi/a screens in human neurons link lysosomal failure to ferroptosis. Nature Neuroscience 24, 1020–1034. 10.1038/s41593-021-00862-0

33. Fearnhead, H.O., Dinsdale, D., and Cohen, G.M. (2000). An interleukin-1 β -converting enzyme - like protease is a common mediator of apoptosis in thymocytes. FEBS Letters 375, 283–288. 10.1016/0014-5793(95)01228-7

34. Yamanaka, K., Saito, Y., Sakiyama, J., Ohuchi, Y., Oseto, F., and Noguchi, N. (2012). A novel fluorescent probe with high sensitivity and selective detection of lipid hydroperoxides in cells. RSC Advances 2, 7894–7900. 10.1039/c2ra20816d

35. Lee, R.G., Rudler, D.L., Raven, S.A., Peng, L., Chopin, A., Moh, E.S.X., McCubbin, T., Siira, S.J., Fagan, S.V., DeBono, N.J., et al. (2023). Quantitative subcellular reconstruction reveals a lipid mediated inter-organelle biogenesis network. Nature Cell Biology 26, 57–71. 10.1038/s41556-023-01297-4

36. Haberkant, P., and Holthuis, J.C.M. (2014). Fat & fabulous: Bifunctional lipids in the spotlight. Biochimica et Biophysica Acta (BBA) - Molecular and Cell Biology of Lipids 1841, 1022–1030. 10.1016/j.bbalip.2014.01.003

37. Sassano, M.L., Tyurina, Y.Y., Diokmetzidou, A., Vervoort, E., Tyurin, V.A., More, S., La Rovere, R., Giordano, F., Bultynck, G., Pavie, B., et al. (2025). Endoplasmic reticulum–mitochondria contacts are prime hotspots of phospholipid peroxidation driving ferroptosis. Nature Cell Biology 27, 902–917. 10.1038/s41556-025-01668-z

38. von Krusenstiern, A.N., Robson, R.N., Qian, N., Qiu, B., Hu, F., Reznik, E., Smith, N., Zandkarimi, F., Estes, V.M., Dupont, M., et al. (2023). Identification of essential sites of lipid peroxidation in ferroptosis. Nature Chemical Biology 19, 719–730. 10.1038/s41589-022-01249-3

39. Drummen, G.P.C., van Liebergen, L.C.M., Op den Kamp, J.A.F., and Post, J.A. (2002). C11-BODIPY581/591, an oxidation-sensitive fluorescent lipid peroxidation probe: (micro)spectroscopic characterization and validation of methodology. Free Radical Biology and Medicine 33, 473–490. 10.1016/s0891-5849(02)00848-1

40. Jelinek, A., Heyder, L., Daude, M., Plessner, M., Krippner, S., Grosse, R., Diederich, W.E., and Culmsee, C. (2018). Mitochondrial rescue prevents glutathione peroxidase-dependent ferroptosis. Free Radical Biology and Medicine 117, 45–57. 10.1016/j.freeradbiomed.2018.01.019

41. López-Doménech, G., and Kittler, J.T. (2023). Mitochondrial regulation of local supply of energy in neurons. Current Opinion in Neurobiology 81, 102747. 10.1016/j.conb.2023.102747

42. Crowley, L.C., Christensen, M.E., and Waterhouse, N.J. (2016). Measuring Mitochondrial Transmembrane Potential by TMRE Staining. Cold Spring Harbor Protocols 2016, pdb.prot087361. 10.1101/pdb.prot087361

43. Venkatraman, K., Lee, C.T., and Budin, I. (2024). Setting the curve: the biophysical properties of lipids in mitochondrial form and function. Journal of Lipid Research 65, 100643. 10.1016/j.jlr.2024.100643

44. Cogliati, S., Enriquez, J.A., and Scorrano, L. (2016). Mitochondrial Cristae: Where Beauty Meets Functionality. Trends in Biochemical Sciences 41, 261–273. 10.1016/j.tibs.2016.01.001

45. Mann, J., Reznik, E., Santer, M., Fongheiser, M.A., Smith, N., Hirschhorn, T., Zandkarimi, F., Soni, R.K., Dafré, A.L., Miranda-Vizuete, A., et al. (2024). Ferroptosis inhibition by oleic acid mitigates iron-overload-induced injury. Cell Chemical Biology 31, 249–264.e247. 10.1016/j.chembiol.2023.10.012

46. Cohen, N., and Sanders, T. (2014). Nematode locomotion: dissecting the neuronal–environmental loop. Current Opinion in Neurobiology 25, 99–106. 10.1016/j.conb.2013.12.003

47. Dixon, S.J., and Olzmann, J.A. (2024). The cell biology of ferroptosis. Nature Reviews Molecular Cell Biology 25, 424–442. 10.1038/s41580-024-00703-5

48. Harayama, T., and Riezman, H. (2018). Understanding the diversity of membrane lipid composition. Nature Reviews Molecular Cell Biology 19, 281–296. 10.1038/nrm.2017.138

49. Kagan, V.E., Mao, G., Qu, F., Angeli, J.P.F., Doll, S., Croix, C.S., Dar, H.H., Liu, B., Tyurin, V.A., Ritov, V.B., et al. (2016). Oxidized arachidonic and adrenic PEs navigate cells to ferroptosis. Nature Chemical Biology 13, 81–90. 10.1038/nchembio.2238

50. Qiu, B., Zandkarimi, F., Bezjian, C.T., Reznik, E., Soni, R.K., Gu, W., Jiang, X., and Stockwell, B.R. (2024). Phospholipids with two polyunsaturated fatty acyl tails promote ferroptosis. Cell 187, 1177–1190.e1118. 10.1016/j.cell.2024.01.030

51. Phadnis, V.V., Snider, J., Varadharajan, V., Ramachandiran, I., Deik, A.A., Lai, Z.W., Kunchok, T., Eaton, E.N., Sebastiany, C., Lyakisheva, A., et al. (2023). MMD collaborates with ACSL4 and MBOAT7 to promote polyunsaturated phosphatidylinositol remodeling and susceptibility to ferroptosis. Cell Reports 42, 113023. 10.1016/j.celrep.2023.113023

52. Fry, M.Y., Navarro, P.P., Hakim, P., Ananda, V.Y., Qin, X., Landoni, J.C., Rath, S., Inde, Z., Lugo, C.M., Luce, B.E., et al. (2024). In situ architecture of Opa1-dependent mitochondrial cristae remodeling. The EMBO Journal 43, 391–413. 10.1038/s44318-024-00027-2

53. Ikon, N., and Ryan, R.O. (2017). Cardiolipin and mitochondrial cristae organization. Biochimica et Biophysica Acta (BBA) - Biomembranes 1859, 1156–1163. 10.1016/j.bbamem.2017.03.013

54. Winnikoff, J.R., Milshteyn, D., Vargas-Urbano, S.J., Pedraza-Joya, M.A., Armando, A.M., Quehenberger, O., Sodt, A., Gillilan, R.E., Dennis, E.A., Lyman, E., et al. (2024). Homeocurvature adaptation of phospholipids to pressure in deep-sea invertebrates. Science 384, 1482–1488. 10.1126/science.adm7607

55. Fernandopulle, M.S., Prestil, R., Grunseich, C., Wang, C., Gan, L., and Ward, M.E. (2018). Transcription Factor–Mediated Differentiation of Human iPSCs into Neurons. Current Protocols in Cell Biology 79, e51. 10.1002/cpcb.51

56. Takeda, H., Matsuzawa, Y., Takeuchi, M., Takahashi, M., Nishida, K., Harayama, T., Todoroki, Y., Shimizu, K., Sakamoto, N., Oka, T., et al. (2024). MS-DIAL 5 multimodal mass spectrometry data mining unveils lipidome complexities. Nature Communications 15, 9903. 10.1038/s41467-024-54137-w

57. Harayama, T., Hashidate-Yoshida, T., Fleuriot, L., Aguilera-Romero, A., Hamano, F., Ri, K., Morimoto, R., Debayle, D., Shimizu, T., and Riezman, H. (2024). A highly efficient gene disruption strategy reveals lipid co-regulatory networks. bioRxiv, 2020.2011.2024.395632. 10.1101/2020.11.24.395632

58. Brinkman, E.K., Chen, T., Amendola, M., and van Steensel, B. (2014). Easy quantitative assessment of genome editing by sequence trace decomposition. Nucleic Acids Research 42, e168–e168. 10.1093/nar/gku936

59. Yusa, K., Zhou, L., Li, M.A., Bradley, A., and Craig, N.L. (2011). A hyperactive piggyBac transposase for mammalian applications. Proceedings of the National Academy of Sciences 108, 1531–1536. 10.1073/pnas.1008322108

60. Dobin, A., Davis, C.A., Schlesinger, F., Drenkow, J., Zaleski, C., Jha, S., Batut, P., Chaisson, M., and Gingeras, T.R. (2013). STAR: ultrafast universal RNA-seq aligner. Bioinformatics 29, 15–21. 10.1093/bioinformatics/bts635

61. Anders, S., Pyl, P.T., and Huber, W. (2015). HTSeq—a Python framework to work with high-throughput sequencing data. Bioinformatics 31, 166–169. 10.1093/bioinformatics/btu638

62. Love, M.I., Huber, W., and Anders, S. (2014). Moderated estimation of fold change and dispersion for RNA-seq data with DESeq2. Genome Biology 15, 550. 10.1186/s13059-014-0550-8

63. Wiśniewski, J.R., Zougman, A., Nagaraj, N., and Mann, M. (2009). Universal sample preparation method for proteome analysis. Nature Methods 6, 359–362. 10.1038/nmeth.1322

64. Rappsilber, J., Mann, M., and Ishihama, Y. (2007). Protocol for micro-purification, enrichment, pre-fractionation and storage of peptides for proteomics using StageTips. Nature Protocols 2, 1896–1906. 10.1038/nprot.2007.261

65. Demichev, V., Messner, C.B., Vernardis, S.I., Lilley, K.S., and Ralser, M. (2019). DIA-NN: neural networks and interference correction enable deep proteome coverage in high throughput. Nature Methods 17, 41–44. 10.1038/s41592-019-0638-x

66. Cox, J., and Mann, M. (2008). MaxQuant enables high peptide identification rates, individualized p.p.b.-range mass accuracies and proteome-wide protein quantification. Nature Biotechnology 26, 1367–1372. 10.1038/nbt.1511

67. Kohler, D., Staniak, M., Tsai, T.-H., Huang, T., Shulman, N., Bernhardt, O.M., MacLean, B.X., Nesvizhskii, A.I., Reiter, L., Sabido, E., et al. (2023). MSstats Version 4.0: Statistical Analyses of Quantitative Mass Spectrometry-Based Proteomic Experiments with Chromatography-Based Quantification at Scale. Journal of Proteome Research 22, 1466–1482. 10.1021/acs.jproteome.2c00834

68. Brenner, S. (1974). The Genetics of Caenorhabditis Elegans. Genetics 77, 71–94. 10.1093/genetics/77.1.71

69. Porta-de-la-Riva, M., Fontrodona, L., Villanueva, A., and Cerón, J. (2012). Basic *Caenorhabditis elegans* Methods: Synchronization and Observation. Journal of Visualized Experiments 64, e4019. 10.3791/4019

70. Vicencio, J., Martínez-Fernández, C., Serrat, X., and Cerón, J. (2019). Efficient Generation of Endogenous Fluorescent Reporters by Nested CRISPR in Caenorhabditis elegans. Genetics 211, 1143–1154. 10.1534/genetics.119.301965

71. Ershov, D., Phan, M.-S., Pylvänäinen, J.W., Rigaud, S.U., Le Blanc, L., Charles-Orszag, A., Conway, J.R.W., Laine, R.F., Roy, N.H., Bonazzi, D., et al. (2022). TrackMate 7: integrating state-of-the-art segmentation algorithms into tracking pipelines. Nature Methods 19, 829–832. 10.1038/s41592-022-01507-1

